# Pump it up: bioelectric stimulation controls tissue shape and size

**DOI:** 10.1101/2022.11.22.517561

**Authors:** Gawoon Shim, Isaac B. Breinyn, Alejandro Martínez-Calvo, Sameeksha, Daniel J. Cohen

**Affiliations:** Department of Mechanical and Aerospace Engineering, Princeton University, Princeton, 08540, NJ, USA; Department of Quantitative and Computational Biology, Princeton University, Princeton, 08540, NJ, USA; Princeton Center for Theoretical Science, Princeton University, Princeton, 08540, NJ, USA; Department of Chemical and Biological Engineering, Princeton University, Princeton, 08540, NJ, USA

**Keywords:** epithelia, bioelectric stimulation, ion transport, hydrostatic pressure, multicellularity

## Abstract

Epithelial tissues sheath many organs, separating ‘outside’ from ‘inside’ and exquisitely regulating ion and water transport electromechanically to maintain homeostatic balance and tissue hydrostatic pressure. While it is increasingly clear that the ionic microenvironment and external electric stimuli can affect epithelial function and behavior, the coupling between electrical perturbation and tissue form remain unclear. We investigated this by combining electrical stimulation with three-dimensional epithelial tissues with hollow ‘lumens’—both kidney cysts and complex intestinal stem cell organoids. Our core finding is that physiological strength electrical stimulation of order 1-3 V/cm (with both direct and alternating currents) can drive powerful and rapid inflation of hollow tissues through a process we call ‘electro-inflation’, inducing up to a threefold increase in tissue volume and striking asymmetries in tissue form. Electro-inflation is primarily driven by field-induced ion crowding on the outer surface of the hollow tissue that creates an ion gradient across the epithelial shell, which drives increased ionic flux mediated by ion channels/transporters and subsequent osmotic water flow into the lumen. This influx generates hydrostatic pressure, and inflation results from a competition between this pressure and cell cytoskeletal tension. We validated these interpretations with computational models connecting ion crowding around tissues to tissue mechanics. Electrically stimulated cysts and organoids also exhibited pronounced asymmetry, where the epithelial shell thickened on the cathode-facing side and thinned on the anode-facing side of the tissue. We discovered that this process is the result of 3D electrotaxis–directed migration of cells in an electric field–causing a redistribution of cells around the shell. The ability of electrical cues to dramatically regulate tissue size and shape highlight the key role of the electrical micro-environment and the potential offered by manipulating these signals.

---

Epithelia—sheet-like multicellular tissues with strong cell-cell adhesion—are foundational to animal life [1] as they can separate ‘outside’ from ‘inside’ by self-assembling into 3D, lumenized (hollow) structures such as tubules (e.g. kidney ducts), intricately folded surfaces (gut epithelium, choroid plexus), and hollow cysts (e.g. lung alveoli, organoids) [2]. Epithelia are intrinsically electromechanical structures and play three critical, interlinked roles in lumenized structures to maintain homeostasis [3]: (1) regulation of endogenous ionic currents via ion transport [4, 5]; (2) regulation of water transport and osmolarity in response to ion gradients (180 L/day in the kidney [6]); and (3) generation and support of hydrostatic pressure resulting from water transport, all of which can be experimentally measured [7–10]. The electromechanical sequence of ion and water transport has been intensively studied, with the general conclusion being that the direction of the transepithelial potential, *not* the membrane polarity, is what ultimately drives water transport, primarily through intercellular junctions [7, 11–15]. Further, the relative permeability and transport in a given epithelium to different ions contribute to the overall magnitude and direction of these potentials and subsequent water transport [13].

Ion transport-mediated pressurization in complex 3D systems such as cysts and branching intestinal organoids has become increasingly recognized as a key factor regulating morphogenesis [7–9] by providing the system biomechanical cues for size control and morphology. Transport across the epithelial layer controls cellular proliferation, differentiation, and rearrangement through luminal depositions of ions and proteins that affect cellular signaling and coordination as well as build-up of hydrostatic pressure that affects cortical tension, tissue stiffness, and maturation of tight junctions in tissues [16]. Given the broad importance of epithelia and their intrinsic electromechanical nature, we investigated how externally manipulating ionic currents affected the growth dynamics of 3D epithelial tissues.

Bioelectric manipulation of epithelia is rooted in the 19th c. discoveries of Matteuci [17] and du Bois Reymond [18] that damage to skin (a stratified epithelium) induces a transient ‘wound current’, which we now know both arises from local disruption of homeostatic ionic currents [19–21] and can occur in any disruptive process spanning healing, embryonic development, and disease progression [19, 20, 22–27]. Crucially, cells sense and respond to these ionic currents through a variety of mechanisms including electrophoretic and electrokinetic displacement of membrane-bound receptors and activation of ion transporters [20, 21, 28, 29]. In 2D epithelia, responses to external electric fields include regulation of cell volume and homeostasis [30], transcriptional changes [21, 31], and ‘electrotaxis’ (galvanotaxis)—the directed migration of cells down ionic gradients (typ. 1 V/cm) [20, 32]. In this case, 2D epithelia undergo particularly dramatic electrotactic collective migration [33–38] which can be precisely controlled by spatiotemporally varying the electric field to induce arbitrary maneuvers or accelerate scratch wound healing [19, 33, 39, 40]. Here, we extend this work into 3D epithelial tissue models (hollow kidney cysts and intestinal organoids) in an attempt to manipulate tissue morphology.

We used the MDCK kidney cell cyst as our initial model system for luminal epithelial structures as MDCKs form single-layered, hollow cysts that are extremely well studied and known to recapitulate key developmental processes including apicobasal polarization of cells, active ionic and water transport, strong cell-cell junctions, and control of cell growth and proliferation [2, 9, 41]. To electrically stimulate these cysts, we grew them within a hydrogel matrix (Geltrex or Matrigel) that we embedded within a modified version of our SCHEEPDOG electrobioreactor [34, 35, 39, 40] that allowed us to combine: live imaging; media perfusion for bioreactor culture; and closedloop electrical stimulation over a range of electric field strengths. Given our prior epithelial electrotaxis data, we hypothesized that we would see strong, 3D electrotaxis, but we instead observed a novel response upon electrically stimulating these cysts. Rather than directional migration, the cysts dramatically and rapidly increased in volume, inflating continuously when we applied an electric field and deflating upon removal of the field. We investigated this and found that the inflation rate varies directly with the magnitude of the electric field. This inflation is mechanically mediated by cortical tension, and we showed through the use of inhibitors that the actomyosin network mechanically stabilizes and constrains growth of the cyst, akin to a spherical pressure vessel. Next, we pharmacologically inhibited apical and basal ion channels/transporters to reveal that cellular ion transport strongly mediates the inflation response. We developed a first-principles computational model for this inflation supporting the concept that external electric bias could ultimately drive increased water transport. Interestingly, we observe from confocal 3D imaging the asymmetric deformation of cysts as they inflate that can be modulated by switching field polarity back and forth, which we attribute as the result of 3D electrotactic migration towards the cathode. Finally, we demonstrate that this ‘electro-inflation’ response generalizes to other lumenized models such as intestinal organoids.

## Results

### Direct current electrical stimulation drives inflation in hollow epithelial cysts

To study how lumenized 3D structures respond to DC electrical stimulation, we stimulated MDCK epithelial cysts with a DC electric field using a modified version of the SCHEEPDOG bioreactor [34, 39] containing two stimulation electrodes, electrochemical safeguards, perfusion lines, and a large, glass coverslide foundation to support high resolution 3D imaging during stimulation (Supp. Fig. 1, Methods). Single MDCK cells were seeded into a basement membrane hydrogel (Geltrex) on top of the coverslip platform and cultured for 96 h as they self-assembled into spherical, lumenized structures approximately 50 *μ*m in diameter (Fig. 1a). The resulting cysts typically had a single hollow lumen, occasionally containing debris from cells having undergone apoptosis during lumen development. Reported field strengths *in vivo* are 𝒪(1 V/cm), but the current density is a key design parameter [20, 33, 42], so we used a current-controlled supply and fixed microfluidic bioreactor dimensions to deliver 0.7 mA/mm^2^ to 1.5 mA/mm^2^ (equivalent to 1V/cm to 4V/cm) to suspended tissues and validated stimulation uniformity using both finite element simulations (Supp. Fig. 2) and real-time channel voltage monitoring. We observed dramatic changes upon stimulation, not via migration as expected, but via rapid volumetric inflation (a 2X increase in crosssection and nearly 3X increase in volume), as shown in 3D confocal sections in Fig. 1b and Supp. Video 1. Closer examination revealed two distinct phenotypes: 1) overall increases in volume as the luminal space expands and 2) asymmetric deformation of the epithelium, with one side thickening while the other thins (Supp. Video 2), and we proceeded to analyze each phenotype in turn.

**Fig. 1.**
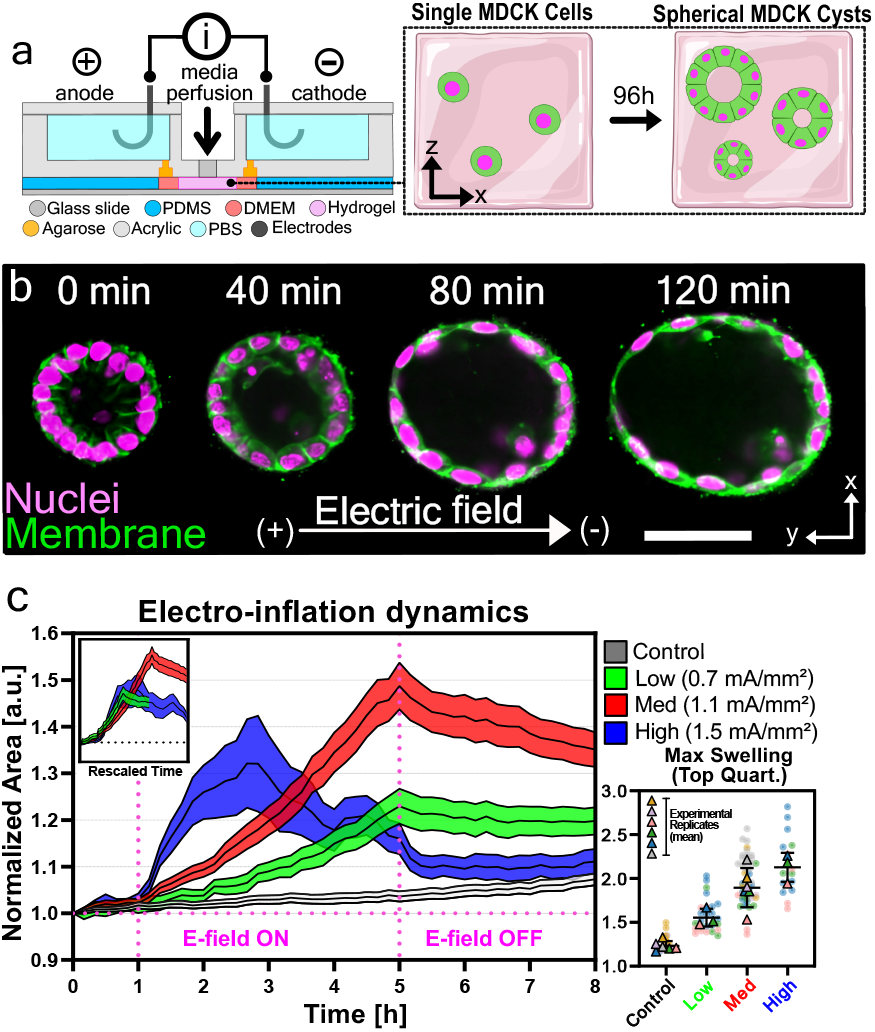
Electrical stimulation increases volume of lumenized epithelia. **(a)** Cross-section of the electro-bioreactor. Inlay: Single MDCK cells form single-layered spherical cysts with hollow lumens after 96 hours of culture. **(b)** Direct current electrical stimulation drives accelerated inflation in MDCK cysts. After 2 hours of stimulation, cysts approximately double in size. **(c)** Inflation dynamics averaged from cysts stimulated with no current (Black; N = 5, n = 144), low current (Green; 0.7 mA/mm^2^, 1 V/cm; N = 3, n = 101), medium current (Red; 1.1 mA/mm^2^, 2.25 V/cm; N = 5, n = 192) and high current (Blue; 1.5 mA/mm^2^, 3.5 V/cm; N = 3, n = 68), normalized to cyst area at *t* = 0 h. Stimulation begins at 1 h and ends at 5 h, as shown by the vertical dotted lines. The asterisk indicates cysts rupture. Shaded error bars show 95% Confidence Interval. Inlay: inflation curves collapse to a master curve after rescaling time using a powerlaw fit (See Methods). Outlay: Top quartile of maximum normalized area for all cysts across conditions. Triangles represent experimental averages while squares represent biological replicates.

To better understand the striking tissue volume change induced by electrical stimulation, we first characterized the broad dynamics of the inflation process and the relationship between input stimulation and output inflation by varying the input field strength over three levels: low (1V/cm; 0.7 mA/mm^2^), medium (2.25 V/cm; 1.1 mA/mm^2^), and high (3.5 V/cm; 1.5 mA/mm^2^). All initial experiments were performed using time-lapse (10 min/frame), phase contrast imaging as follows: control observation for 1 h; electrical stimulation for 4 h; relaxation without the field for 3 h. Data were analyzed by segmenting the phase contrast images at each timepoint and calculating the crosssectional area of the cysts at the spherical equator using a custom ImageJ script (Methods). The crosssectional area for each individual cyst was normalized to the value at *t* = 0 and filtered for artifacts (filter pipeline presented in Supp. Fig. 3 along with all raw data). Each epithelial cyst originates from a single cell, meaning cysts are clonal and subject to any heterogeneity in the population, and cysts themselves also exhibit considerable heterogeneity in culture [41, 43]. To quantify how the temporal dynamics of the inflation response varied with electric field magnitude, we directly plotted the complete dataset of mean cross-sectional area of all cysts per field strength over time with the 95-percent confidence interval in Fig. 1c (Supp. Video 3; the complete set of all raw traces is shown in Supp. Fig. 3). In all cases, cysts began to inflate within 20 minutes of electrical stimulation, exhibiting a clear and direct relation between the rate and extent of inflation and applied electric field strength. In all cases, stimulated tissues exhibited a pronounced difference in inflation relative to unstimulated tissues (Fig. 1c, gray). In the strongest field case, inflation proceeded so rapidly that many cysts appeared to burst, likely from being over-pressurized (Fig. 1c, blue, ‘*’). These burst cysts were capable of repairing themselves but the majority ceased inflation post-collapse. More generally, we found that peak inflation volumes appeared capped and could not be increased indefinitely with field strength or duration. In both types of inflation, with or without rupture, we did not observe cellular death over the course of our experiments (Supp. Video 4, 5: EthD1 dead-cell detection assay). Prolonging the duration of electrical stimulation at medium current also did not inflate the system further (Supp. Fig. 4). These data together suggest that there is a mechanical limit to how much pressure the cyst can withhold, possibly set by the cytoskeletal structure within the cyst, a hypothesis we explore in the subsequent section. Once electrical stimulation was removed, the cysts gradually decreased in volume until they converged on the volume of time-matched control cysts, surprisingly (Fig. 1c from 5+ hrs; Supp. Fig. 5 for longer duration). To explore the generality of electro-inflation dynamics, we re-scaled all inflation curves using a power-law fit (see methods, Supp. Fig. 6), after which the data collapsed onto a single master curve (Fig. 1c, inlay). This shows that the kinetics of electro-inflation are self-similar across experimental conditions, and the mechanisms governing the response are conserved across stimulus strength. This is further emphasized by the independence of the cysts’ relative change in cross-sectional area on their initial area (Supp. Fig. 7), supporting the claim that this process is general and agnostic of the intrinsic properties of the cyst. Given this, we used a ‘medium’ electric field strength (1.1 mA/mm^2^, 2.25 V/cm) for all further experiments, which we calculated corresponds to an inflation rate of approximately 2 pL/min if we assume cysts are spheres and generalize from the equatorial cross-section changes plotted in Fig. 1c.

We next compared the relative potency of electro-inflation to contemporary, pharmaceutical-based approaches to inducing inflation that target ion channels/transporters [44–47] and osmotic flow [48]. Here we evaluated the effect of the commonly used swelling drug, forskolin, which acts by elevating cyclic AMP (cAMP) levels to trigger downstream protein kinase A (PKA) and the apical chloride channel cystic fibrosis transmembrane conductance regulator (CFTR) (Supp. Fig. 8) [49, 50]. Interestingly, we found that electrically mediated inflation far outstrips forskolin-induced inflation over the experimental duration, with forskolin reported to primarily drive inflation over a multi-day period [51]. Overall, these findings suggest that electrical stimulation activates a powerful and reversible inflation mechanism in the epithelial cysts, raising the question of how this is regulated.

### Electrically-actuated ion transport and water flow drive inflation

We next investigated how electrical stimulation facilitates inflation in epithelial cysts. As electrically-induced inflation happens on a timescale of several hours, we ruled out the possibility that proliferation is the major drive behind the initial volume increase, although it may play a role in the long-term relaxation dynamics where we observed a bimodal response where some cysts remain significantly inflated while others gradually collapsed (Supp. Fig. 5). Hence, we hypothesized that there are two possible mechanisms for electrical inflation: (1) direct actuation of water influx into the lumen through aquaporins; or (2) actuation of ion influx, which would create electro-osmotic gradients to drive paracellular water influx.

The accepted mechanism of luminal volume increase consists of two stages. First, ion influx alters the electrical potential of the luminal space, which triggers electrically-driven influx of ions of opposing charge to achieve electrical balance. Second, this net influx of salt ions into the lumen creates a strong osmotic gradient that drives water into the lumen via both paracellular and trans-cellular routes [52–54]. Fig. 2a presents a cartoon of what this process might look like in the case of electro-inflating cysts [49] where the apical and basal surfaces might both contribute to ion flow. We aimed to inhibit both ion influx from the cell into the lumen by targeting apical ion channels and from the extracellular environment into the cell by targeting basal ion channels/transporters, to test if field-induced ion transport regulated cyst inflation.

**Fig. 2.**
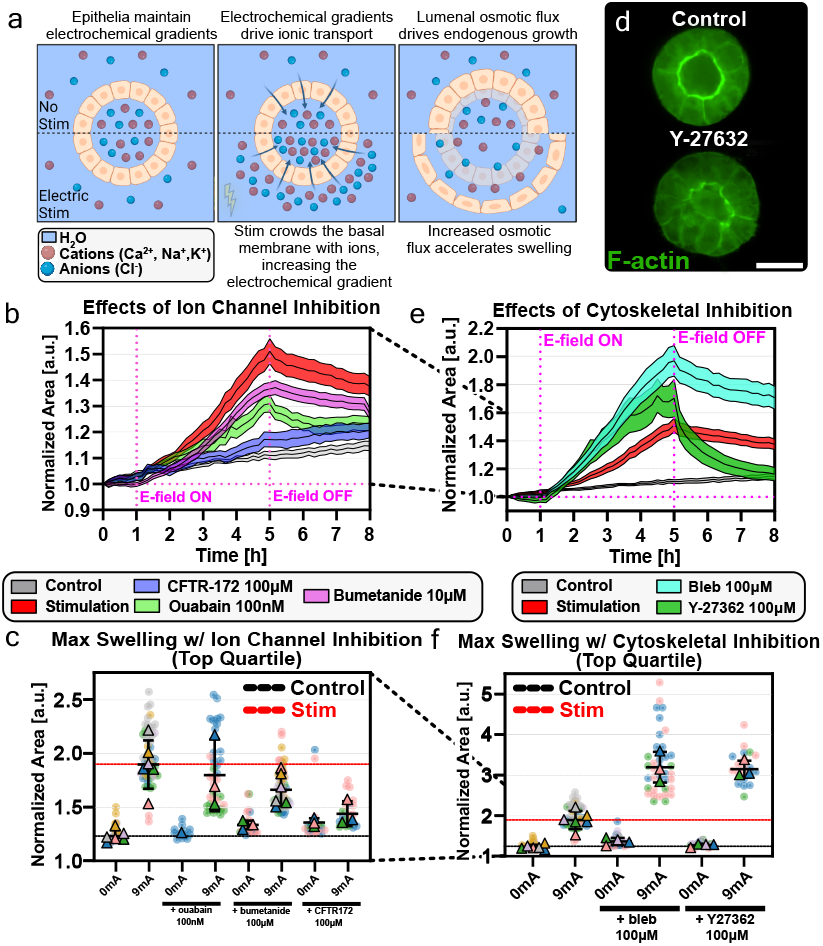
Ion transport and cytoskeletal mechanics mediate electrical stimulation-mediated inflation. **(a)** Schematic of the process by which MDCK cysts endogenously inflate, and how the inflation mechanism is modulated by electrical stimulation. **(b)** Inflation dynamics averaged for cysts with no electrical stimulation (Black; N = 5, n = 150), stimulation with no inhibitors (Red; N = 5, n = 150), and stimulation with 50*μ*M (Pink; N = 3, n = 101) and 100*μ*M (Blue; N = 3, n = 80) CFTR-172 treatment, and 100 nM Ouabain treatment (Green; N = 3, n = 149). **(c)** SuperPlot of the maximum normalized area of all cysts treated with apicobasal ion channel/transporter inhibitors, with and without electrical stimulation. Colored squares represent individual cysts, with the correspondingly colored triangle representing the experimental average. Violin mean lines represent the average of all experiments within that condition. See Supp. Fig. 11 for replicates and statistical tests. **(d)**, (Top) Equatorial confocal slice of a cyst stained for F-actin, showing the actomyosin cortex that makes up the apical domain of the structure. (Bottom) a cyst that has been treated with Y-27632, a ROCK inhibitor, with disrupted cytoskeletal network. **(e)** Inflation dynamics averaged for cysts with no electrical stimulation (Black; N = 5, n = 150), stimulation with no inhibitors (Red; N = 5, n = 150), and stimulation with Y-27632 (Teel; N = 3, n = 84) and blebbistatin (Green; N = 3, n = 147) treatment. **(f)** SuperPlot of the maximum normalized area of all cysts treated with cytoskeletal-related inhibitors, with and without electrical stimulation. See Supp. Fig. 11 for replicates and statistical tests. Shaded error bars in b and e represent 95% Confidence Interval.

First, we used viability and permeability assays to confirm that the concentration of pharmacological inhibitors we used for our assays did not damage the epithelial barrier function nor induced cytotoxicity (Supp. Fig. 10). We then electrically stimulated MDCK cysts that we had treated with various inhibitors. These data are presented in ‘SuperPlot’ format [55] in Fig. 2c and all statistical tests are presented in Supp. Fig. 11. We first targeted the role of Na+/K+ ATPase, a basal ion pump that transports sodium, potassium, and chloride into cells. As inhibition of the pump using ouabain in renal cysts was reported to reduce lumen size and abolish fluid secretion[9, 56, 57], we hypothesized that inhibiting Na+/K+ ATPase would also decrease the magnitude of electro-inflation. We indeed observed a strong reduction in luminal volume increase in ouabain-treated cysts, suggesting that disrupting ion transport into the epithelia heavily affects ion secretion into the lumen. This is consistent with recent data implicating electrical stimulation of this ATPase as being useful for improved outcomes during kidney transplantation[58]. We further tested basal ion transport by targeting the charge-balanced Na-K-Cl cotransporter (NKCC) with the inhibitor bumetanide, diluted to 100*μ*M as used in previous studies on transepithelial transport in MDCK cysts [49, 59], and we also saw significant reduction in the electro-inflation response.

We next assessed chloride transport into the lumen by targeting the cystic fibrosis transmembrane conductance regulator (CFTR), an apical (lumen-facing) chloride channel known to regulate chemical-induced inflation of MDCK cysts and other epithelial lumenized structures [9, 49, 53]. Using a CFTR inhibitor, CFTR(inh)-172 [49, 60], dramatically reduced electro-inflation (Fig. 2b). Both CFTR(inh)-172 and bumetanide required a higher concentration (50*μ*M and 100*μ*M) than the amount used for MDCK monolayer assays to achieve a noticeable effect in 3D (Supp. Fig. 12), although we observed no cytotoxic effects. While these concentrations have also been validated in numerous recent 3D organoid studies[8, 61], we do note that higher levels of CFTR(inh)-172 can potentially affect volume-sensitive outwardly rectifying chloride channels (VSORCC) in kidney tissues, meaning the effects we observe here may implicate chloride transport beyond CFTR itself [62]. Either way, these data demonstrate that electro-inflation is strongly mediated by ion transport into the lumen via apical and basal ion channels/transporters that lead to osmotic water transport and hydrostatic pressure changes.

### Cystokeletal mechanics constrain inflation

There is clearly a mechanical component to electrically-induced tissue inflation, and the situation is akin to a spherical pressure vessel where hydrostatic pressure is balanced by hoop stresses in the vessel walls. Here, the mechanical balance between luminal pressure and epithelial stress appears to define a physical limit to the maximum size MDCK cysts can inflate to based on the plateaus evident at 5h in Fig. 1c, and the early rupture that occurs with overly rapid inflation (e.g. 1.5 mA/mm^2^ or 3.5V/cm stimulation). We hypothesized that these plateaus and rupture can both be linked to cortical cytoskeletal effects in the cysts limiting strain beyond a certain point in a viscoelastic fashion.

Considering the cyst as a simple pressure vessel, we predicted that reducing actomyosin contractility would effectively relax the pre-stress in the cyst shell and allow MDCK cysts to inflate to a greater extent. Using blebbistatin, a myosin inhibitor, and Y-27632, a Rho kinase (ROCK) inhibitor, we inhibited actomyosin contractility of the cysts previous to electrical stimulation (Supp. Video 6). Our data showed that the actomyosin inhibitor treatment successfully disrupted the cortical actomyosin network in MDCK cysts (Fig. 2d) and significantly increased both the inflation rate and the maximum volume that inflated cysts could reach. Both drugs promoted increased inflation, with blebbistatin treatment showing the strongest effect–up to 1.5-fold higher than positive controls, which is in keeping with data showing blebbistatin, and myosin II inhibition, more effectively softens cellular stiffness relative to ROCK inhibition via Y-compound [63] (Fig. 2e, 2f).

These data further show that molecular transport into the lumen and cytoskeletal mechanics are intimately coupled in facilitating and regulating the electro-inflation response, respectively. To fully capture the biophysical mechanisms behind this phenomenon, we moved towards developing a model in which we couple electrokinetics with established 1D elastic models for pressurized epithelial structures.

### A mechanical model for MDCK cysts inflation in an electrolyte solution under an externally applied electric field

Given the importance of tissue hydrostatic pressure, there has been a surge of interest in physical models that can describe tissue electromechanics related to pressure [16, 46, 64–71]. We build on these studies here to develop an interactive model that allows us to relate an external electric field to inflation of the lumen. At a bulk level, our data resemble hydraulic pumping into a spherical pressure vessel, which prompted us to develop a continuum model to capture our experimental observations and test our inflation hypotheses *in silico* to help determine the primary driving forces.

We first hypothesize that an externally applied electric field is able to increase the electrolyte ion concentration around the basal surface of the cyst [72, 73]. This occurs because the charged outer surface of the cyst attracts counter-ions via Coulomb force, creating an ionic cloud the screens the surface charges, usually referred to as the diffuse double layer (DDL) [72, 73], and whose ion concentration increases with external field strength. Such ion crowding would lead to permeation of ions into the lumen, as the concentration in the lumen is lower than in the DDL. As ion concentration increases in the lumen, a sufficiently strong osmotic pressure gradient between the lumen and the DDL establishes, able to drive water into the lumen which results in cyst inflation (see Fig. 2a and Fig. 3a) [46, 67, 68, 70]. As field strength increases, we argue that cyst inflation is enhanced as such gradients become stronger.

**Fig. 3.**
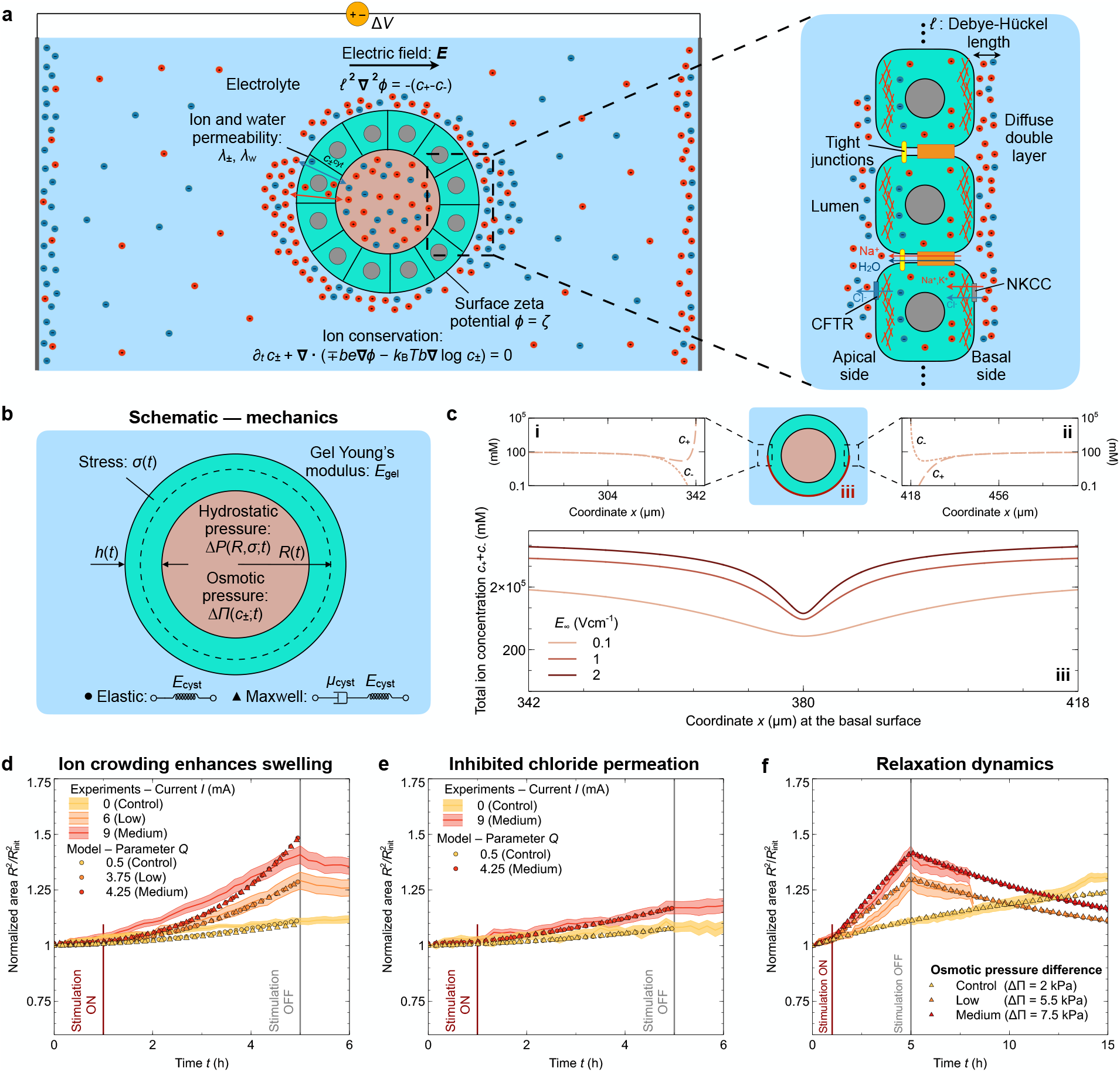
Viscoelastic model of MDCK cysts under electrical stimulation recapitulates the inflation dynamics. **a**, Schematic of the 1D and 2D models, in which we consider a spherical MDCK cyst embedded in an electrolyte. An external electric field is applied at *t* = 0^+^. The rightmost panel displays a zoom-in schematic of the epithelium showing the different ion and water transport mechanisms considered in the 1D model, as well as a magnified of the electric double cloud of ions surrounding the basal surface of the MDCK cyst (not to scale). **b**, Schematic of the mechanics used in the 1D model, in which we couple hydrostatic pressure Δ*P*, osmotic pressure ΔΠ (which depends on the electrokinetics shown in panel **a**, gel elasticity, and stress *σ* of either an elastic (circles) or viscoelastic (triangles) spherical shell of mean radius *R* and thickness *h*. **c**, Panels **i** and **ii** show the steady-state concentration of positive *c*_+_ and negative *c*_−_ ion species in the DDL at the basal surface, both at the anode-facing **i** and cathode-facing **ii** poles of the cyst. Panel **iii** displays the total ion concentration *c*_+_ + *c*_−_ along the basal surface for different values of the far-field electric field ***E***_∞_. **d**-**f**, Normalized area as a function of time obtained from experiments (curves) and 1D model (circles and triangles), for different electric field strengths (see Supp. Info.). Panel **e** corresponds to the knock-out experiment inhibiting ion transport through the epithelium. **f**, Same as in panels c and d but using a simplified 1D model, in which the osmotic pressure increases due to stimulation but remains constant over time (see Supp. Info.).

where *ε* is the permittivity of the electrolyte, *e* is the elementary charge, ***r*** and *t* denote the positional vector field and time, respectively, and ***v***_*±*_ is the velocity of ion species, i.e., ***v***_*±*_ = ∓ *be***∇** *ϕ* −*k*_B_*Tb* **∇**ln *c*_*±*_. The first term accounts for electrostatic forcing of ions, and the second term takes into account diffusion of ions, where *b* = *D/*(*k*_B_*T*) is the mobility of ions and *D* the diffusivity constant of ions in the outer medium, *k*_B_ is the Boltzmann constant, and *T* is the temperature. At the basal surface, we impose a uniform zeta potential *ζ* and zero ion flux.

To obtain the distribution of ions at the basal surface when an uniaxial electric field is externally applied, we numerically solve Eqs. (**??**) in time until steady state is reached (see theory Supp. Info.). Fig. 3c shows the ion concentration of positive *c*_+_ and negative *c*_−_ ions at different locations of the basal surface. Panels c(i) and c(ii) show the concentration of positive and negative ions as a function of the distance from the basal surface, at the left and right side of the cyst, respectively. As expected, we find a dipolar distribution of ion charges due to diffusion and electrophoretic effects [72]. Panel c(iii) displays the total ion concentration *c*_+_ + *c*_−_ as a function of the *x* coordinate along the basal surface for different field strengths, i.e., *E*_∞_ = 0.1, 1, and 2 V cm^−1^. We find that the concentration of ions at the basal surface increases as the strength of the externally applied field increases, as we hypothesized and as suggested by a simple scaling analysis: *c*∼ *εE*_∞_*/*(*e*ℓ_c_), where ℓ_c_ is the characteristic length scale of the DDL. Thus, ion crowding at the basal domain of the cyst, which increases with applied field strength, enhances the transport of ions into the lumen, due to both osmosis and an increased trans-epithelial potential.

To further test if ion crowding induced by an external electric field can drive cyst inflation, we considered a simplified version of the model. In particular, we assumed a uniform ion concentration along the basal surface and considered an effective increase of ion concentration due to the external field, which is controlled by a dimensionless parameter *Q*, proportional to the surface charge of the basal domain and the external field (see theory Supp. Info.). We also assumed a cyst is either an elastic or a viscoelastic spherical shell embedded in an electrolyte solution. Initially, we consider the shell behaves as a linearly elastic material and impose cell volume conservation, considering that cells do not proliferate during acute cyst inflation (see theory Supp. Info.). Additionally, cyst mechanics, hydrostatic and osmotic pressure, and lumen volume conservation are coupled to the applied electric field via electrokinetics at the DDL and transport of ions across the epithelium (see sketches in Figs. 3a and b). The main ingredients of this model are well validated in other approaches and contexts involving lumenized structures, which we built on by newly introducing electromechanical coupling to an external electric field [7, 16, 64, 66–71, 74, 75] (see theory Supp. Info. for more details). In particular, the main driving mechanism underlying inflation is ion transport due a strong gradient between the external ion crowd at the basal surface and the lumen, which is controlled by *Q*, followed by water transport from the electrolyte into the lumen due to osmotic pressure differences, which is balanced by hydrostatic pressure in our model.

We first focus on the control case with no external electric field. The positively charged basal (outer) surface accumulates a screening layer of counter-ions, i.e., the DDL. Such ion crowding works to neutralize the surface charge but also increases the concentration of ions around the cyst and triggers the transport of species into the lumen due to osmotic effects, as the concentration of ions in the lumen is lower than in the DDL. Moreover, the electric potential across the epithelium also increases due to ion crowding [72, 73], which drives the transport of ions across the epithelium, creating a strong osmotic imbalance between the lumen and the basal surface, that drives water across the epithelium leading to cyst inflation [67] (Figs. 3d-f, yellow curves).

When an external electric field is applied, ion crowding over the basal (outer) surface increases asymmetrically (see Fig. 3c and Supp. Info.). Such increase in ion concentration at the basal surface triggers increased ion transport into the lumen, in turn driving enhanced osmotic transport of water and subsequent rapid inflation. Our minimal model recapitulates the inflation dynamics of MDCK cysts both in the absence of an external electric field and during electrical stimulation, as displayed in Fig. 3d. Moreover, our model is also able to capture the MDCK cyst inflation dynamics when ion transport is inhibited (Fig. 3e), as well as the long term relaxation behavior of cysts after stimulation is removed (see Fig. 3f, Supp. Fig. 13, and theory Supp. Info.). While our model well captures inflation dynamics, building on our work to fully resolve cyst inflation and ion distribution at the basal surface, epithelium, and lumen, is an important direction for future work. We hope our work inspires new models that incorporate such additional complexities present in experiments.

Moreover, this approach allowed us to explore different material models; of particular interest given both the rate-dependent rupture shown with the highest current density we tested, and the fact that cysts gradually relaxed back to its initial volume after stimulation was removed. Rate dependent material processes can imply viscoelasticity, so we considered the two simplest models to incorporate viscoelasticity at the spherical shell force balance: Maxwell and Kelvin-Voigt materials. A Maxwell material behaves as an elastic solid at times shorter than the so-called Maxwell time, which is the ratio between viscosity and Young’s modulus *μ*_cyst_*/E*_cyst_, and exhibits a viscous resistance to deformation at times larger than the Maxwell time. A Kelvin-Voigt material behaves oppositely. Here, we find that the Maxwell model was effectively identical to the purely elastic model (Figs. 3d-f), while we were unable to fit a Kelvin-Voigt material to our inflation dynamics (see theory Supp. Info.). Together, these data suggest that viscous effects take place at time scales longer than the actual inflation event. Additionally, we find that both linear elastic and Maxwell materials coupled to electro-kinetics yield a maximum hydrostatic pressure of 1.2-1.8 kPa, for low and medium stimulation, respectively. While we cannot directly measure these values, they are consistent with the magnitudes reported in previous measurements [10, 16].

### Electrically-directed cell migration causes tissue asymmetry

In addition to inflation itself, we also observed pronounced and unexpected asymmetry developing in inflating cysts. Visualizing this symmetry breaking required confocal sectioning, which consistently revealed that the thickness of the hemisphere of the cyst facing the anode (‘left’ in all of our figures) thinned over the stimulation period, while the cyst wall on the cathode-facing side tended to thicken (Fig. 4a). The thinning side (anode-facing) collapsed down to such an extent that the nuclei re-aligned and stretched tangent to the surface, akin to a squamous epithelium. Conversely, cells on the thickening side (cathode-facing) appeared to crowd together and elongate like a columnar epithelium, with nuclei orienting perpendicular to the surface. Changing the field polarity changed the direction of thickening and the asymmetry disappeared when electrical stimulation was stopped, meaning that this phenomenon was both reversible and dependent on the field direction (Supp. Video 7). The asymmetry also seemed to be independent of the inflation response, as we could observe strong asymmetry even in weakly inflating cysts (Supp. Video

**Fig. 4.**
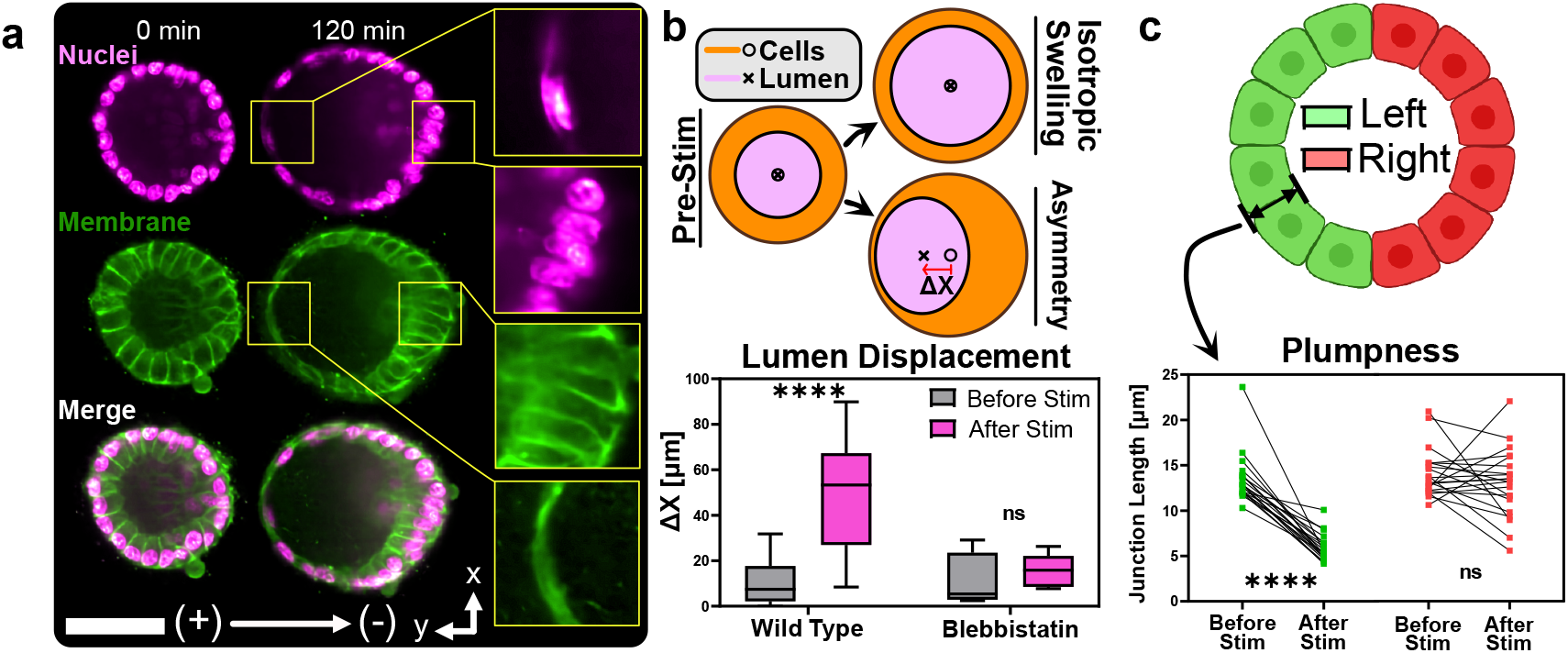
Cyst symmetry is broken by electrical stimulation. **(a)** DC stimulation causes asymmetrical thickening of the cathodefacing half of MDCK cysts. Inlay: Opposing halves of the cyst experience nuclear re-orientation and junctional thinning or thickening. **(b)** Lumen displacement as measured by the difference between basal and apical membrane centroid position. Cytoskeletal inhibition with Blebbistatin almost entirely abolishes the asymmetry response. **(c)** Junction length averaged over the left and right half of the cyst before and after stimulation (see cartoon). Outlay: The heterotypic inflation response is captured by pairwise comparisons of averaged junction lengths before and at the end of stim (N = 3, n = 376/383). P values are calculated using pairwise parametric unpaired t-tests with Welch’s correction. **** corresponds to P < 0.0001.

To understand the origins of electrical symmetry breaking, we quantified different aspects of the asymmetric response. First, we measured the extent of the asymmetry by collecting confocal data at the equator of each cyst and measuring the displacement of the centroid of the hollow lumen relative to the centroid of the total cyst cross-section, as illustrated in the cartoon and plot in Fig. 4c. We found that electrical stimulation caused the lumen to displace significantly towards the anode to create structural asymmetry. We next quantified thickness changes in the epithelial shell by measuring the cell-cell junction geometry at the cyst midplane both before electrical stimulation and at the time of maximal inflation and asymmetry. Binning cysts into ‘left’ (anode-facing) and ‘right’ (cathode-facing) hemispheres broadly revealed that junctions on the left side always thinned by 2X relative to their initial lengths, while junctions on the right side exhibited a more heterogeneous response, spanning mild thinning to moderate thickening (Fig. 4c, Supp. Fig. 14). These effects are captured in greater detail by analyzing junction length and orientation as a function of junction angular position before stimulation and at the time of maximal inflation (Supp. Fig. 15, 16), where the data emphasize a linear relationship between junction orientation and angular position that can be modulated by electrical stimulation.

Structural asymmetry clearly results from a mechanical effect, which we initially hypothesized was due to asymmetric cortical tension, perhaps from apical constriction. However, when we inhibited actomyosin contractility with blebbistatin under confocal imaging, we still observed a weak asymmetry (Supp. Video 6) suggesting that pure contractility cannot explain the full effect. We next investigated whether electro-kinetic shear forces resulting from the electric field might be deforming the gel matrix or directly shearing the surface of the tissue [76]. Our analysis of stimulated gels in the absence of cells did not reveal any biased deformation pattern, negating gel deformation as a mechanism (Supp. Fig. 17). More telling was when we tested an alternate tissue model– human induced pluripotent stem cells (hiPSC) hollow spheroids[77, 78]. These spheroids were similar in morphology to MDCK cysts but, crucially, exhibited asymmetry in the opposite polarity of MDCK cysts–that is they thinned on the right (cathode-facing) side (Supp. Video 9). This effect rules out a simple physical force explanation and suggested a biological basis for asymmetry. We also noted that although hiPSC spheroids exhibited asymmetry, they did not inflate under electrical stimulation (Supp. Fig. 18), further supporting our claim that luminal inflation and thinning-thickening asymmetry are independent phenomena.

A parsimonious biological explanation was simply that electrical stimulation induced canonical cathodal electrotaxis (directed cell migration in an electric field), except on the surface of a hollow sphere. As MDCK cells are known to electrotax towards the cathode[33, 37–39], 3D electrotaxis could explain cells crowding on the cathode-facing side of the cyst and migrating away from (stretching) the anode-facing side. To confirm that the 3D electric field had an appropriate geometry for electrotaxis, we performed a numerical simulation of a uniaxial electric field wrapping around a spherical tissue (modeled as a pure dielectric here). This simulation (Fig. 5b) revealed that the spherical cyst geometry should experience a unique field well suited for cathode-directed electrotaxis. The computed field predicts divergent migration on the ‘left’ pole, which would account for thinning, convergent migration on the ‘right’ pole, which would account for compression and thickening, and strong tangent electrotaxis at the two poles orthogonal to the bulk field (Fig. 5c). We validated the prediction by tracking individual cell nuclei during electrical stimulation, which revealed a stereotyped pattern of migration that largely followed the predicted field lines (Fig. 5c; Supp. Video 10), strongly suggesting electrotaxis to be the cause of asymmetry. As a final confirmation, we pharmacologically reversed electrotaxis using LY294002, a phosphoinositide 3-kinase (PI3K) inhibitor that is known to reverse the electrotactic migration direction of keratocyte monolayers [79, 80]. After validating that PI3K inhibition reversed 2D electrotaxis direction in MDCK cells (see Methods and Supp. Video 11), we inhibited PI3K in MDCK cysts and observed a striking reversal of asymmetry polarity and apparent electrotaxis towards the anode (Fig. 5d, Supp. Video 12). These data are further supported by our previous results showing that blebbistatin reduces asymmetry, as blebbistatin treatment for MDCK monolayers is known to reduce directional electrotaxis [37]. Finally, the ability to interactively reverse asymmetry by switching field polarity is consistent with our prior demonstrations of reversing collective migration in 2D electrotaxis[33]. Overall, these data strongly support that electrotaxis around a 3D shell can create asymmetric geometries by altering cellular crowding and epithelial shell thickness.

Both our computational model of ion crowding and water transport and established literature on epithelial transport [13, 14, 81] suggest that the direction of water transport should not depend on which surface of the epithelium is exposed to the electric field, depending instead on the transepithelial potential itself. To confirm this, we electrically stimulated

**Fig. 5.**
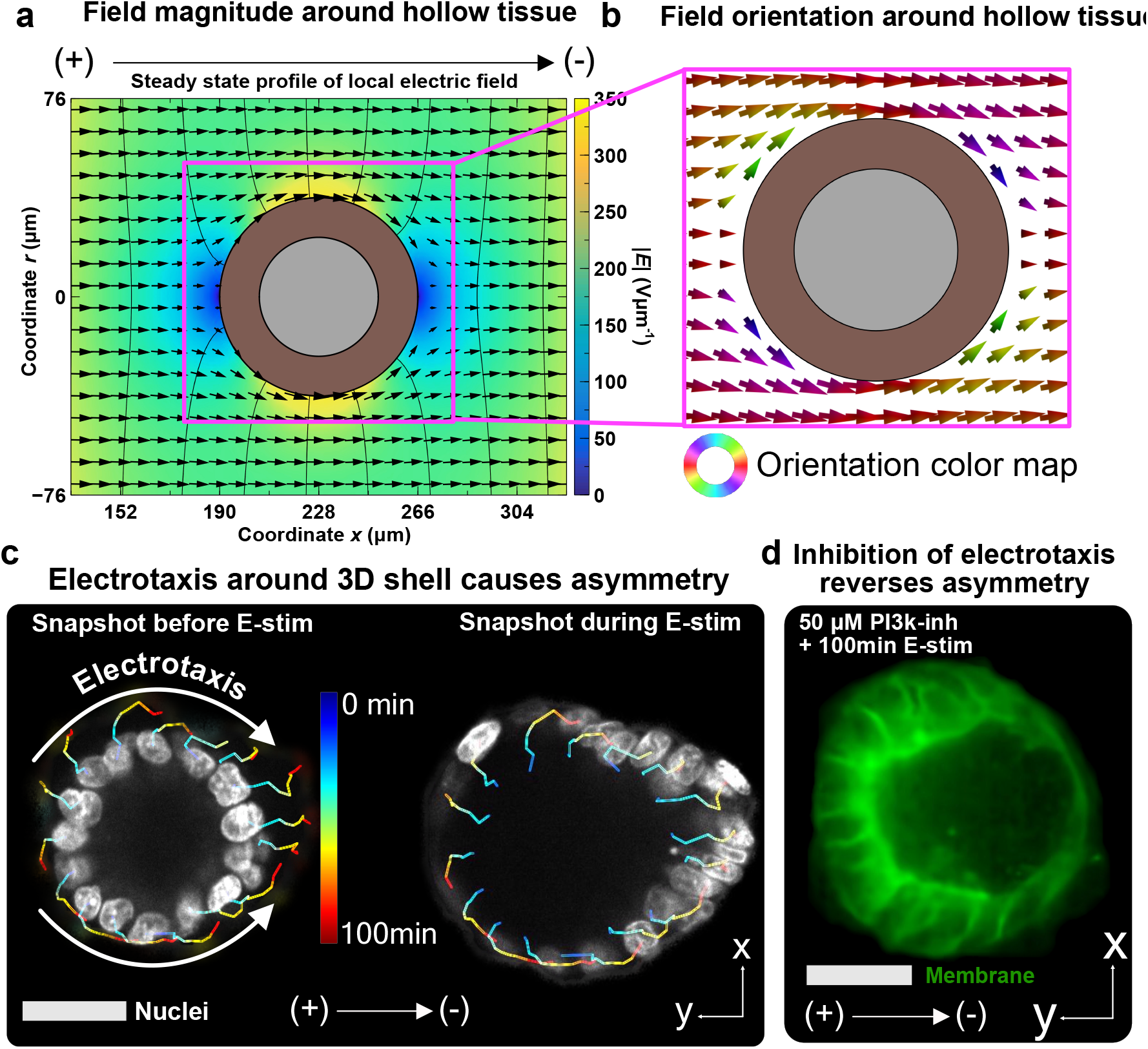
Structural asymmetry under electrical stimulation is driven by electrotaxis. **(a)** Heatmap of 2D axisymmetric electrical potential *ϕ* isolines (black curves) and electric field (arrows and color) outside a spherical shell fixed in space representing a lumenized structure. **(b)** Electric field vectors local to the lumenized structure, colored by their orientation. **(c)** Nuclear tracks of an MDCK cyst breaking symmetry after 100min of electric stimulation. Cell motion, while restrained to the surface of the cyst, follows the electric field shown in panels **(a)** and **(b). (d)** Inhibition of the primary electrotactic pathway PI3k entirely reverses cyst asymmetry in identical electric fields, implicating electrotaxis as the driver of asymmetry. All scale bars are 50 *μ*m.

**Fig. 6.**
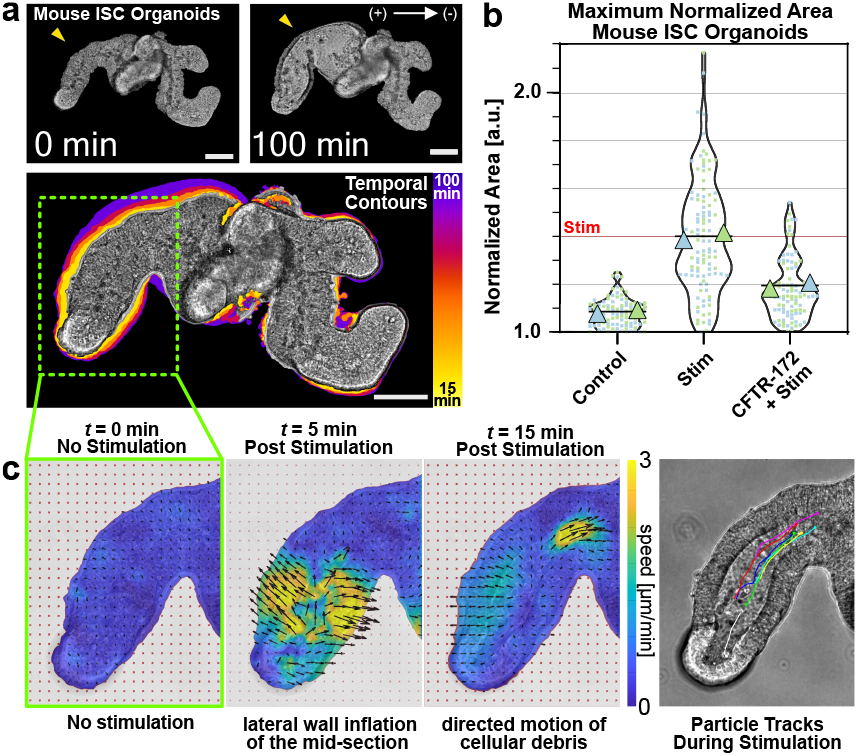
Electrical stimulation-induced inflation in mouse ISC organoids. **(a)** (Top) Mouse intestinal organoid before (*t* = 0 min) and after (*t* = 100 min) electrical stimulation at medium current. (Bottom) Overlay of outlines for each timepoint. **(b)** Maximum normalized area for mouse intestinal organoids with no stimulation (N = 2, n = 103), with 2.5 h electrical stimulation at medium current (N = 2, n = 103), and with 2.5 h electrical stimulation at medium current with 100*μ* CFTR(inh)-172 (N = 2, n = 92). **(c)** (Panels 1-3) Velocity fields of fluid flow in the lumen of electro-inflating mouse intestinal organoids. (Panel 4) Particle tracks of debris in the lumen of electro-inflating mouse intestinal organoids. All scale bars = 100 *μ*m.

MDCK domes grown on a 2D substrate [82], a well-documented pseudo-3D model system. As MDCK domes are inverted luminal structures with an inward-facing basal membrane (opposite that of our cysts), electrical stimulation of dome cultures would directly stimulate the apical, rather than basal, surface. If our model was accurate, electrical stimulation of domes would also cause the luminal space to inflate. Surprisingly, we initially observed electrically induced deflation of MDCK domes (Supp. Video 13). We hypothesized that this deflation could be due to mechanical failure caused by high interstitial stresses at the dome-monolayer boundary during 2D electrotaxis of cells surrounding the domes [82]. To eliminate such electrotactic effects, we once again used the PI3k inhibitor LY294002, which we found could inhibit electrotaxis at lower concentrations that that needed to reverse it. By preventing electrotaxis and eliminating any stresses caused by it, we were able to see that MDCK domes inflated when electrically stimulated in regions of the monolayer exhibiting minimal electrotaxis (Supp. Fig. 19, Supp. Video 14). This confirmed that electrically driven inflation is independent of cell polarity, consistent with the proposed ion crowding mechanism and in agreement with prior findings.

### Generality of electrically driven inflation

The ability to electrically control epithelial hydrostatic pressure has exciting implications for developmental biology, biofabrication, and regenerative medicine if it generalizes [10]. As our data thus far focused on the MDCK (kidney) model, we next investigated how general the electro-inflation mechanism might be for other lumenized epithelia with more complex cell type populations and architectures. As our key test case, we chose the increasingly popular intestinal organoid model derived from mouse intestinal stem cells. Intestinal organoids are complex, lumenized tissues exhibiting multiple cell types, complex 3D morphologies, and recapitulating key aspects of gut function. Moreover, they represent an entirely distinct organ group relative to our MDCK kidney model, while still being known to exhibit spontaneous, CFTR-mediated inflation and collapse dynamics during cell differentiation and budding morphogenesis, making them a good target model [9, 83]. We cultured intestinal organoids using a Matrigel substrate [84] and otherwise identical stimulation conditions to our prior ‘medium’ stimuli (see Methods).

We immediately noticed that intestinal organoids also exhibited electro-inflation, where certain individual lobes or the central cavity inflated quite strongly in many cases (Fig. 6a, Supp. Video 15, 16). Using the CFTR inhibitor CFTR(inh)-172, we proved that CFTR also mediates intestinal organoid electro-inflation, indicating that electrically actuated ion transport may be a general mechanism for the external control of lumenized epithelia and tissue hydrostatic pressure (Fig. 6a). Interestingly, we found significant variability in the response of complex 3D organoid structures, with inflation occurring more often in the main cavity of the organoid compared to the high curvature stem cell crypts, and with instances of directional flow of luminal debris during electro-inflation (Supp. Video 15). We investigated this further and used particle-image-velocimetry (PIV) to visualize both the spatiotemporal dynamics of organoid deformation and the emergence of directional flow. These data are shown in Fig. 4a and highlight three things that we also observed in other samples. First, the crypts (high curvature tips of the organoid lobes) exhibit little overall change in cross-section (compare Fig. 4c, *t* = 0 and *t* = 5 min). Second, the majority of the initial deformation seems to occur along the long axis of a lobe (Fig. 4c, t=5 min). Third, there is clear directional flow of debris that emerges during this process, and Fig. 4c at *t* = 15 min shows a fast moving region of debris that is clearly visible forcing the initial constriction of the lobe open. Particle tracking (Fig. 4c, right) confirmed the majority of debris travel persistently up the lobe, consistent with fluid primarily influxing from the crypt itself. These data emphasize that structural differences in a tissue can contribute to differences in the electro-inflation response, and suggest three hypotheses for future testing. First, crypt cells are known to be mechanically stiffer than the surrounding villi cells, implying crypt zones would deform less under hydrostatic forces [85]. Alternately, perhaps there is differential expression of ion transporters within different cell types in the organoid, which would alter influx patterns. Finally, and more broadly, the complex 3D geometries might induce ionic ‘shadowing’ similar to the asymmetric ion distributions described earlier in our 2D model, which might explain the observed heterogeneous response.

## Discussion

Our work demonstrates how epithelia are electrically controllable pumps, where external, DC electric fields of physiological magnitudes 𝒪(1V/cm) control 3D tissue size and shape (symmetry breaking) through the combined effects of hydrostatic pressure regulation and electrically guided collective cell migration (electrotaxis). Here, external electrical stimulation of 3D tissues drives tissue pressurization and extreme inflation by actively controlling the existing electromechanical infrastructure of ion crowding/transport and paracellular water transport. In parallel, spherical tissues break geometric symmetry and dramatically thin and thicken on opposite poles in response to electrically driven, 3D collective cell migration (electrotaxis) around the surface of the tissue. These findings emphasize both the importance of the bioelectric microenvironment on tissue form and function, and the opportunity to harness this effect to regulate 3D tissue form and function.

We characterized the direct relationship between applied electric field strength and the rate, and extent, of inflation. Broadly, stronger electric field strengths induced faster, more powerful swelling. However, overly strong fields induced premature rupture due to rapid inflation. The inflation process itself is clearly linked to external electric fields triggering ion-channel mediated ion transport and subsequent paracellular water transport into the lumen of the tissues to balance the osmotic gradient, causing net inflation. We found that ion channels/transporters that maintain transepithelial homeostasis by controlling cellular ion transport were primary regulators of electro-inflation, and their disruption reduced the electro-inflation response down to 1/3. The net build-up in hydrostatic pressure has mechanical consequences and inflation is constrained by cytoskeletal components. In particular, relaxing actomyosin contractility significantly enhanced inflation ((Fig. 4d, e, f)) indicating the reinforcing and constraining role of the cytoskeleton on inflation. This cytoskeletal role, along with the rate-dependent rupture during rapid inflation suggests a viscoelastic component that deserves further study.

Interestingly, the effect of electric fields on tissue shape seemed decoupled from the inflation process itself, instead depending on the local field geometry around the tissue to drive ‘electrotaxis’. While 2D collective electrotaxis sees tissue sheets literally translating across planar culture dishes, 3D electrotaxis here played out on a closed, 3D surface, resulting in unique boundary conditions that caused cells to congregate at one pole (a sink) while trying to actively depart from the other pole (a source), leading to a spatial difference in both cell density and shell thickness.

That environmental electric fields can alter morphogenetic processes raises intriguing possibilities for developmental biology and engineering, especially as water transport and tissue pressure are now known to contribute to development and tissue homeostasis [70]. A classic example of the role of hydrostatic pressure regulated by ion transport and the electricalmicroenvrionment is Kupffer’s vesicle that forms during zebrafish embryogenesis. The vesicle begins without a lumen and then gradually pressurizes over time and contributes to left-right developmental axis asymmetry in a collective migration process around a lumenized space [46, 59, 86]. Similarly, the mammalian amniotic sac both lumenizes and develops distinct cell asymmetry analogous to what we observe here; playing a key role in downstream embryogenesis[87, 88]. Lastly, kidney homeostasis fundamentally depends on maintenance of physiological ion gradients. This behavior is both a direct function of the electrical-microenvironment, and can be modulated by external electric fields to better preserve kidney function during transplantation procedures [58].

The sheer speed of electro-inflation allows much faster perturbation and control than traditional pharmacological agents. Similarly, the efficiency, throughput, and relatively inexpensive infrastructure complement delicate mechanical methods (e.g. micro-pipette aspiration and AFM). Further, these benefits play out in a fully 3D context without requiring dimension reduction to 2D, allowing arbitrary 3D tissue structures to be perturbed. Future work to more precisely localize field stimulation could significantly enhance the level of precision here as well.

If we think of electro-inflation as an actuation process, then the epithelium becomes an incredibly active material–a living, conformal surface with controllable fluid transport, and a number of interesting bioengineering applications become worth exploring, such as in bio-soft robotics [7, 89–91]. Here, hydraulic actuation offers an unexpected and potentially useful alternative to traditional muscle-based actuators. Such engineering applications require a design framework, so it is promising that our preliminary computational models successfully capture and predict the key responses because this could help with formal design and optimization efforts. Overall, it is exciting to consider new uses of bioelectric control of multicellular systems in areas spanning tissue engineering, developmental biology, biosensing, and even water purification [92].

## Materials and Methods

### Cell Maintenance and Culture

MDCK-II wild type canine kidney cells and RFP ECadherin MDCK-II cells (courtesy of the Nelson Lab, Stanford University) were cultured in low-glucose (1 g/L) DMEM with phenol red (Gibco, USA; powder stock), 1 g/L sodium bicarbonate, 1% streptomycin/penicillin (Gibco, USA), and 10% fetal bovine serum (Atlanta Biological, USA)–DMEM++. Cells were incubated at 37°C and 5% CO2 in humidified air. ACTB mGFP (beta actin) human induced pluripotent stem cells (hiPSCs) were purchased from the Allen Institute for Cell Science and cultured in mTeSR1 media (Stemcell Technologies, Vancouver, Canada) supplemented with 10 *μ*M Y-27632 (Stemcell Technologies, Vancouver, Canada) for the first 24h of culture, then without for subsequent days.

### MDCK Cyst Culture

To ensure that the hydrogel containing MDCK cysts was within the stimulation zone and to control their shapes and sizes, a silicone stencil of 250 *μ*m thickness (Bisco HT-6240, Stockwell Elastomers) containing two 7.5 × 10mm square microwells was cut and applied to the center of the glass culture substrate using a razor writer (Cameo, Silhouette, Inc.). MDCK-II wildtype cells were suspended in a DMEM++ solution of 2.5e5 cells/mL using a Cytosmart Cell Counter for counting. Gibco Geltrex LDEV-Free Reduced Growth Factor Basement Membrane Matrix (Fisher Scientific) was mixed at a 1:1 ratio with the DMEM++ cell solution. A first layer of Geltrex/DMEM++ solution of 15*μ*L was deposited onto each 7.5×10 mm2 rectangular stencil, then allowed to solidify for 30 min at 37C. 12.5uL of Geltrex/ cell + DMEM++ solution was deposited on top of the first layer, then allowed to solidify for 1 h before the gel was covered with DMEM++ and cultured for 96h.

### Intestinal Stem Cell (ISC) organoid Culture

Mouse intestinal stem cell (ISC) organoids were isolated and harvested from HaloYAP-positive mice (courtesy of the Posfai Laboratory, Princeton University) according to Stemcell Technologies™’s mouse intestinal crypt isolation protocol, then maintained in culture using reagents and protocol from the IntestiCult OGM Mouse Kit (Stemcell Technologies, Vancouver, Canada). Intestinal organoids were disassociated and resuspended into Corning® Matrigel® Growth Factor Reduced (GFR) Basement Membrane Matrix, Phenol Red-free, LDEV-free, 10 mL (Corning) at a 1:1 ratio with organoid growth medium. 27.5*μ*L of the mixture was deposited onto each of the rectangular stencils and allowed to solidify for 15 min at 37C before the gel was covered with organoid growth medium and cultured for 72h.

### Human Induced Pluripotent Stem Cells (hiPSC) Spheroid Culture

hiPSCs were detached from the tissue culture dish surface using accutase (Stemcell Technologies, Vancouver, Canada) on the third day of culture and resuspended into mTeSR1 solution of 3 × 10^6^ cells/mL. The mTeSR1 cell solution was mixed at a 1:1 ratio with Corning® Matrigel® Growth Factor Reduced (GFR) Basement Membrane Matrix, Phenol Red-free, LDEV-free, 10 mL (Corning). 27.5*μ*L of the Matrigel/mTeSR1 cell solution was deposited onto each of the rectangular silicone stencils and allowed to solidify for 15 min at 37C. The gel was then covered with mTeSR1 for 48h.

### MDCK Dome Culture

MDCK-II cells were seeded onto tissue-culture plastic surfaces at 4 × 10^6^ cells/mL density. The tissue culture plastic substrate was first treated with collagen IV dissolved to 50 *μ*g/mL in DI water, which was applied to the dish for 30 min at 37°C, then washed three times with DI water to provide a matrix for cellular adhesion. With the two 7.5 × 10mm square microwell stencil in place, seeding solution of cells was prepared at a density of 2.5 ×10^6^ cells, counted using a Cytosmart Cell Counter (Corning) and 10 *μ*L of the cell solution was pipetted into the stencils. The cells were allowed to settle for 1 h with extra humidification media was added to the periphery of the tissue culture dish. Once the cells had adhered to the substrate, sufficient media was added to fill the dish. The stencils were removed 48 h after incubation for assembly.

### MDCK Monolayer Electrotaxis

MDCK monolayer electrotaxis was performed similarly to recent studies from the lab, and more detailed methods can be found in the respective publications [33–35, 39, 40]. Briefly, 5×5mm MDCK-II monolayers were seeded at a density of 2.5 × 10^6^ cells/mL (10 *μ*L) and given 18h to reach confluency in an incubator kept at 37C and 5% CO_2_. Tissue barriers were pulled before the electric bioreactor was assembled, and tissues were given a 1h control period before stimulated at 9mA for four hours.

### Inhibitor Assays

For actomyosin inhibition, blebbistatin (Selleck Chemical) and Y-27632 (Selleck Chemical) were used at 100*μ*M concentration and added 1h before stimulation. For aquaporin 3 (AQP3) inhibition, copper sulfate (CuSO4, Sigma Aldrich) was used at 12.5*μ*M concentration and added 24h before stimulation. For Na+/K+ ATPase inhibition, ouabain octahydrate (Sigma Aldrich) was used at 100nM and added 24h before stimulation. For CFTR channel inhibition, CFTR(inh)-172 (Cayman) was used at 50*μ*M and 100*μ*M concentration and added 24h before stimulation for MDCK cysts and 48h before stimulation for ISC organoids. For NKCC inhibition, bumetanide was used at 100*μ*M concentration and added 1h before stimulation. For PI3K inhibition, LY294002 (Medchemexpress) was used at 25 *μ*M to stall electrotaxis at and 50 *μ*M to reverse electrotaxis for both cysts and monolayers. The LY294002 treated samples were incubated at 37C for 60 min before being assembled into the bioreactor. Media with the same concentration of LY294002 was used for perfusion. Inhibitors were applied after the patterning stencils were removed to prevent the inhibitors from being absorbed into the PDMS. Identical concentration of inhibitors was supplemented in the perfusion media to prevent the inhibitors from washing out during stimulation.

### Cellular Death and Epithelial Integrity Assays

To test whether our experimental conditions were causing cellular death or loss of epithelial integrity in MDCK cysts, we respectively used Ethidium Homodimer-1 (EthD-1; LIVE/DEAD Viability/Cytotoxicity Kit, Invitrogen) and 4kDa fluorescein Isothiocyanate-Dextran (Sigma Aldrich). Cyst cultures were treated with 1*μ*M EthD-1 solution for 1 h before electrical stimulation to ensure cellular loading. When used with perfusion, an identical concentration of EthD-1 was supplemented in the perfusion media to prevent the reagent from washing out during stimulation. Dextran was used at a concentration of 0.5mg/mL dissolved in DMEM++ solution and added immediately at the start of experiment.

### Electrical stimulation

The uniaxial stimulation device is a modified version of our SCHEEPDOG bioreactor with similar methods from our previous work [34, 35, 39]. A Keithley source meter (Kiethly 2450 Textronix) provided current to the silver and silver chloride electrode pair while a USB oscilloscope (Analog Discovery 2, Digilent Inc.) measured the voltage across a pair of recording electrodes (titanium wire, 0.5 mm diameter, Alfa Aesar) as probes. A custom MATLAB script was written to maintain a constant current density within the microfluidic chamber. Samples were imaged for 1h without any electrical stimulation, 4h with the designated electric field, then 3h without electrical stimulation. A current density of 0.7mA/mm2 corresponds to a field strength of 1V/cm, 1.1mA/mm2 to 2.25V/cm, and 1.5mA/mm2 to 3.5V/cm.

### Timelapse Microscopy

Phase contrast images were acquired on an automated inverted microscope (Zeiss Observer Z1; 3i Marianas System) equipped with an XY motorized stage, Photometrics Prime sCMOS, and Slidebook capture software (Intelligent Imaging Innovations, 3i). The microscope was fully incubated at 37°C, and 5% CO2 was constantly bubbled into media reservoir during perfusion. Imaging for MDCK cysts was performed using a Zeiss 10X/0.30 Plan-Neofluor objective and images were captured every 10 minutes at 100ms exposure. Imaging for ISC organoids was performed using a Zeiss 20X/0.80 Plan-Apochromat objective and images were captured every 1 minute at 100ms exposure. Additional imaging of organoids was performed on an incubated Nikon Ti2 with a Qi2 CMOS camera and NIS Elements software.

Confocal microscopy was performed at the Princeton University Molecular Biology Confocal Imaging Core using a Nikon Ti2 inverted microscope with a Yokogawa W1 spinning disk and SoRa module. Laser lines 405nm, 488nm, and 561nm were used for nuclei imaging with Hoescht 33342, membrane imaging with Memglow 488, and Ecadherin-RFP markers, respectively. 3*μ*m thick optical slices were captured at 200ms exposure with 15% laser power on all channels. Images were captured using a 40x/1.25 silicone oil immersion objective with a Hamamatsu BT Fusion sCMOS (6.5 *μ*m/px) every 10 minutes. The stage was incubated at 37°C and 5% CO2 was constantly bubbled into media reservoir during perfusion. The NL5 fast scanning confocal was used for…

### Fluorescence staining

For nuclear staining, Hoescht 33342 10mg/mL solution (Invitrogen) was diluted to 1:1000. For membrane staining, Membrane dye (Memglow 488, Cytoskeleton Inc.) was diluted to 100 *μ*M concentration. Both dyes were diluted in serum-free media and added to samples 30 min previous to imaging. For fluorescence actin imaging, MDCK cysts embedded in Geltrex/DMEM++ mixture was fixed using 2% glutaraldehyde solution for 15 minutes and washed twice in PBS. The cells were then permeabilized with 0.1% Triton X-100 (Sigma Aldrich) solution in PBS for 10 minutes. cells were washed with PBS-Triton solution twice, then incubated with 1% Bovine Serum Albumin solution (Thermo Fisher) for 15 minutes for blocking. Phalloidin (Alexa 488) was diluted to 1:500 in PBS was added and incubated for 1h, then samples were washed and submerged in PBS for imaging.

### Inflation Quantification

FIJI (https://imagej.net/software/fiji) was used to find the mid-plane of cysts and binarize the images for further analyses. First, the “find focused slices” plug-in was used to find the mid-plane of each cyst within the z-stack. Once all focused slices within the timelapse were found, images underwent a series of morphological functions including “Enhance Contrast”, “Subtract Background’, ‘Find Edges’, and “Gaussian Blur” before thresholding and converting the image to a mask using “Convert to Mask”. In order to fill any breaks across the boundary of the mask as well as fill holes within the interior of the mask, the image was dilated three times before running “Fill Holes”, after which the image was eroded three times. Finally, “Analyze Particles” was used to remove all but the largest object in the image. Binarized masks were imported to MATLAB (Mathworks, 2020b) for quantification, where the regionprops() function was used to calculate the area of each mask over time. These area dynamics are then normalized by the initial area and filtered by their maximal area and rate of inflation to remove cysts that do not inflate. Inflation dynamics are then averaged together within different experimental conditions (stimulation strength, inhibitor, control, etc.). To better showcase the upper limit capabilities of electro-inflation, we sometimes show the top quartile of inflating cysts. To do this, we simply calculate percentile limits for all cysts’ maximal normalized cross sectional area, then filter for cysts above the 75th percentile (or, top quartile).

### Power Law Fitting and Rescaling

To calculate the rescaling factor of the individual Normalized Area (NA) curves, we defined a time *τ* at which NA(*τ*) = *α*, where *α* was chosen to be an arbitrary normalized area. This calculation was repeated for several different choices of *α*, all yielding the same collapse. After calculating *τ* for each experimental condition (current density, *j*), *τ* (*j*) was plotted in log-log and fitted to *τ* ∝ *j*^*μ*^ using linear regression. Finally, the time vector for each individual curve was rescaled as Δ*t* = *t/j*_*μ*_.

### Junction length quantification

Cell junction lengths in the cysts were measured manually using the built-in MATLAB function ginput(). Equatorial confocal slices of the cyst at the beginning and end of stimulation were loaded into MATLAB and opened into a GUI, wherein the user defines the center of the cyst before sequentially selecting the apical and basal ends of each junction in an arbitrary order. Once the endpoints of each junction are saved, a simple script calculates the junction length, orientation, and angular position within the cyst. Junction metrics are then binned by position for plotting.

### Single-cell tracking analysis

Nuclei images from confocal imaging were analyzed using TrackMate plugin in ImageJ. Laplacian of Gaussian (LoG) filter to detect the nuclei diameter. The simple Linear Assignment Particle (LAP) tracker was used to track the nuclei and produce a trajectory over the course of the timelapse video.

### Lumen displacement quantification

Ellipses were fit to both the basal and apical membranes of the cyst at timepoints corresponding to the start of stimulation and the time of maximal swelling/asymmetry. Geometric properties of those ellipses are then loaded into Matlab and the displacement between centroids is calculated. Here, we present only the displacement along the axis of the electric field for ease of visualization.

### Organoid dynamics

Organoid projections were created by z-projecting a time-series as a ‘sum’ projection in FIJI and then switching to the ‘Fire’ look-up-table. Particle-ImageVelocimetry was performed using PIVLab 2.56 in MATLAB [93]. First, image sequences were registered and masked in FIJI and then imported in PIVLab. 2-pass FFT deformation analysis was used with 64/32 pixels for the passes with a 50 percent step-over and scaling performed within PIVLab. To better capture the directed particle flow, we averaged 3 frames (3 minutes) of PIV vector fields to reduce random background noise. Manual tracking in FIJI allowed us to generate trajectories of dead cells and debris within the organoid.

## Supp. Info

All theoretical modeling is discussed in the additional supplementary file. Please refer to Journal-level guidance for any specific requirements.

## Data Availability

All study data are included in the article and/or SI Appendix. Microscopy images and data used to generate figures have been deposited in Zenodo (10.5281/zenodo.7348438).

## Code Availability

Custom codes for image analysis and numerical simulations can be found on GitHub (github.com/CohenLabPrinceton/CystSwelling). Further code is available from the corresponding authors upon reasonable request.

## Acknowledgments

D.J.C. and the project overall were supported by an NIH MIRA Award (R35-GM133574) and an NSF CAREER Award (2046977). G.S. was supported by the Brit and Eli Harari Fellowship. I.B.B. was supported by an NSF GRFP Fellowship. A.M.-C. acknowledges support from the Princeton Center for Theoretical Science and the Human Frontier Science Program through the grant LT000035/2021-C. It is a pleasure to acknowledge enlightening discussions with Profs. Howard A. Stone (Princeton University), and Sean Sun (Johns Hopkins University).

## Author contribution

G.S. and D.J.C. conceived the project. G.S., I.B.B., and D.J.C. designed the research. G.S., I.B.B., and S. performed the experiments. I.B.B. and A.M.-C. performed the theoretical calculations. A.M.-C. performed the numerical simulations. G.S., I.B.B., and D.J.C. performed the image analysis. G.S., I.B.B., A.M.-C., and D.J.C. wrote and edited the manuscript/prepared the figures.

