## Supplement for "Pump it up: bioelectric stimulation controls tissue shape and size"

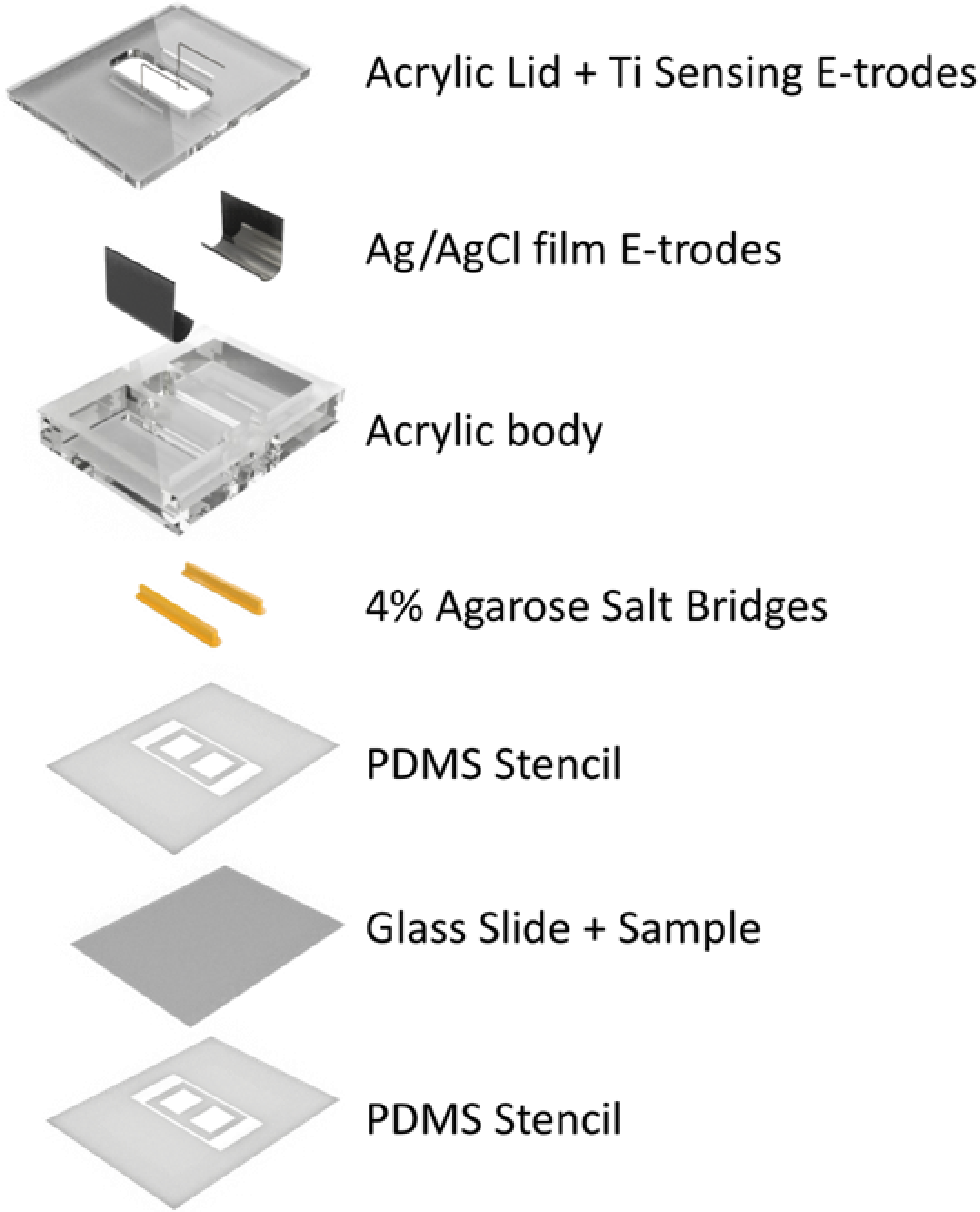


**Supplementary Figure 1** Individual components of the electric bioreactor setup. Silver/silver-chloride electrodes connected directly to a current supply were submerged into a PBS reservoir on either side of the PDMS microfluidics chamber to deliver electric field through agarose salt bridges.

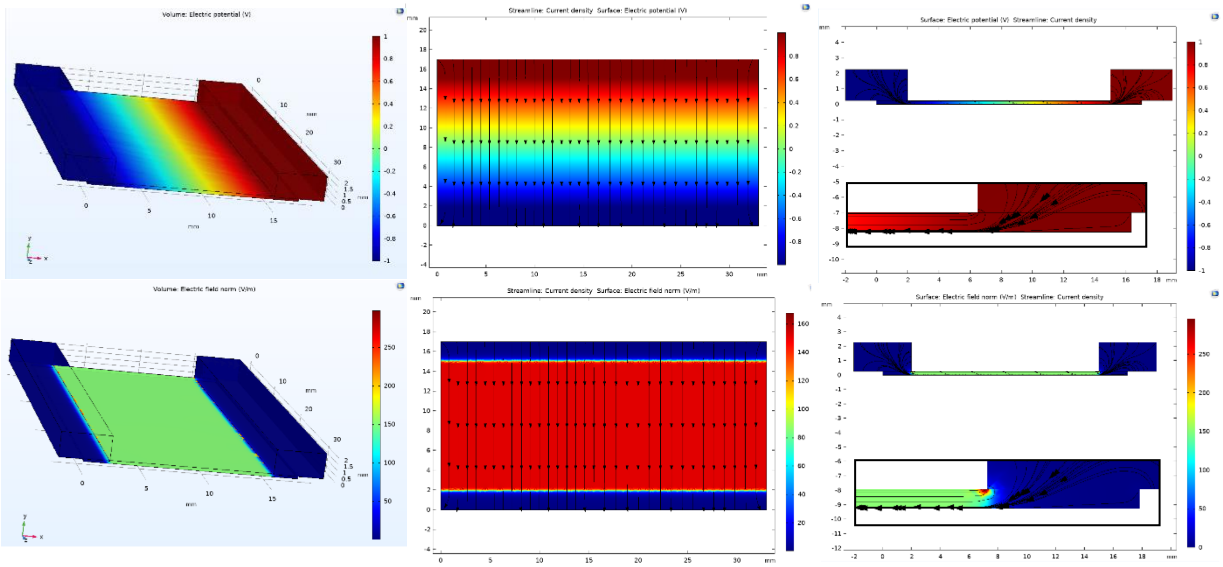


**Supplementary Figure 2 Left**: 3D Finite Element simulation of Electric Potential (*Top*) and Field Strength (*Bottom*) in the bioreactor used for Cyst stimulation. **Middle**: Top-down view of current density (black streamlines) within the microfluidic channel where cysts are cultured and stimulated. **Right**: Cross-sectional side view of current density (black streamlines) within the microfluidic channel where cysts are cultured and stimulated. (**Inset**) Current density at the salt bridge-electrolyte interface shows slight fringing far from the region of the device containing the cysts.

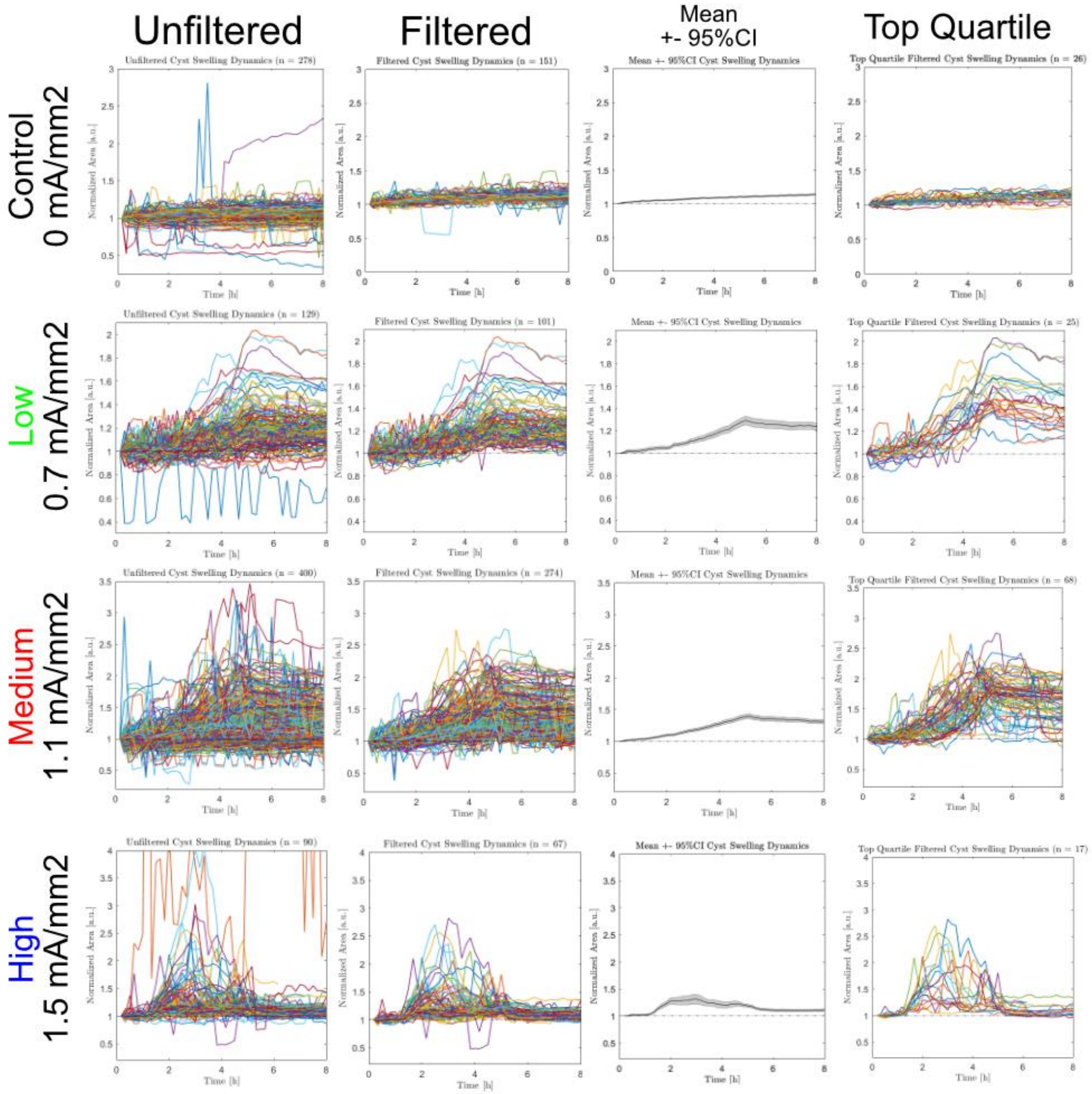


**Supplementary Figure 3** Raw, filtered, averaged, and top quartile (left to right) plots for control, low, medium, and high stimulation (top to bottom). For all experiments, all cysts were binarized to produce raw area plots throughout the duration of the experiment. The raw cyst area plots were filtered for irregularities that arise from image processing, then averaged and plotted alongside the 95% confidence interval. The top quartile was calculated to produce the final plot.

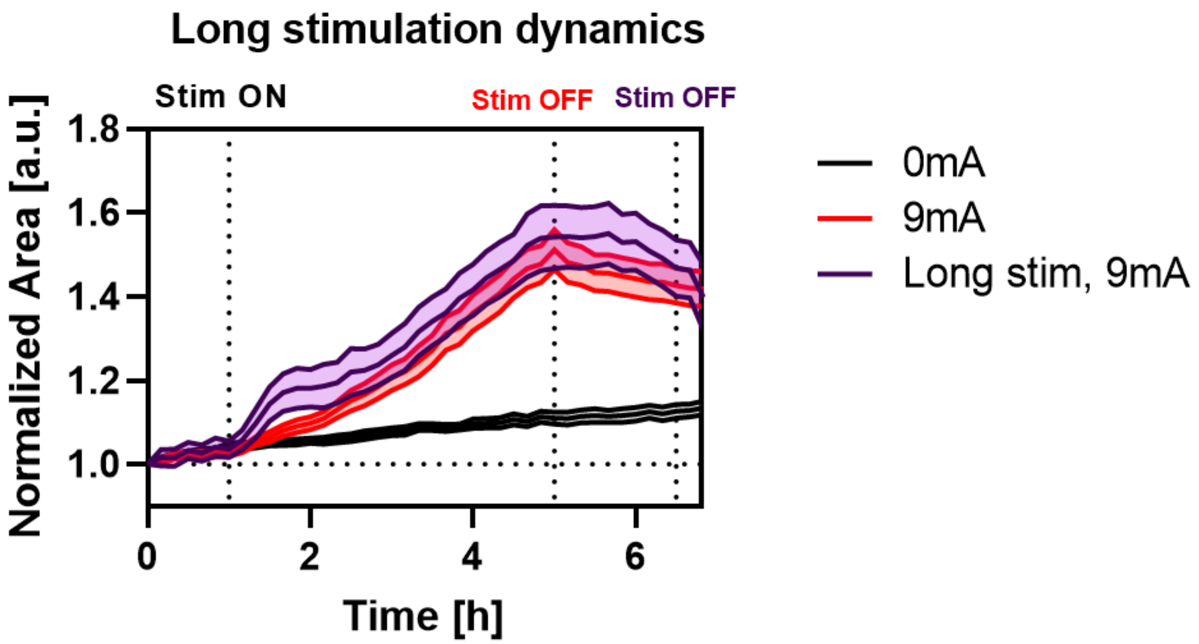


**Supplementary Figure 4** Comparison between MDCK acini stimulated at medium current for 4h (red; stimulation on from 1h – 5h; N = 150) and for 5.5h (purple; stimulation on from 1h – 6.5h; N = 1, n = 72). Longer stimulation does not increase the cyst volume further, suggesting that the maximum inflation magnitude under electrical stimulation is capped for MDCK cysts.

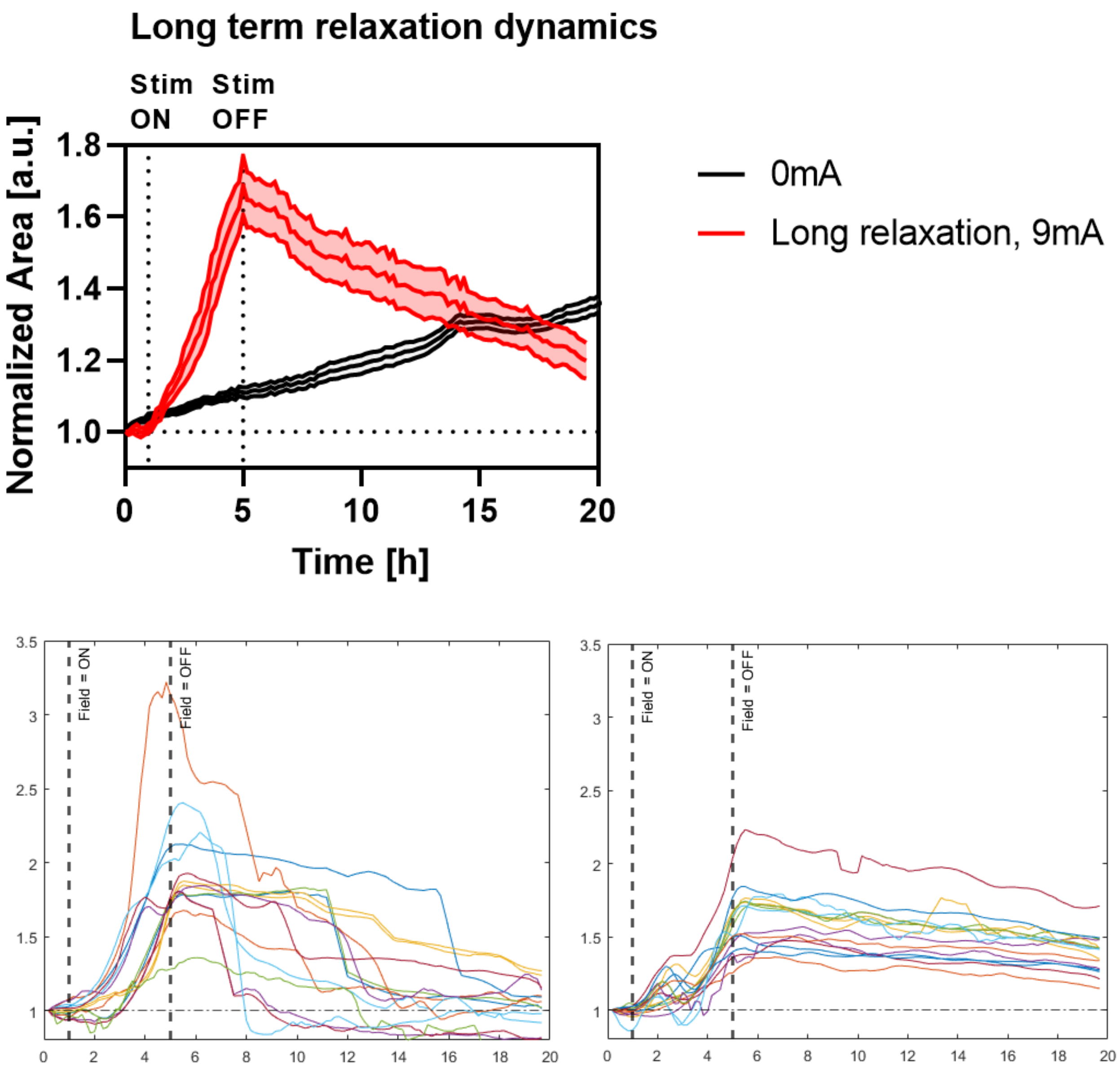


**Supplementary Figure 5** Normalized area as a function of time capturing deflation dynamics of MDCK acini electrically stimulated at medium current stimulation. Stimulation is applied at the 1h mark, maintained for 4h, then removed (N = 1, n = 80). Deflation is slow and gradual. Area dynamics during long relaxation is bimodal. **Top**: Averaged normalized area plot of all cysts electrically stimulated then relaxed (green) and cysts with no electrical stimulation (grey) **Bottom left:** Individual cysts' normalized area plots with rapid deflation, where relaxing cysts suddenly drop in volume. **Bottom right:** Individual cysts' normalized area plots with steady deflation, where they drop in volume slowly.

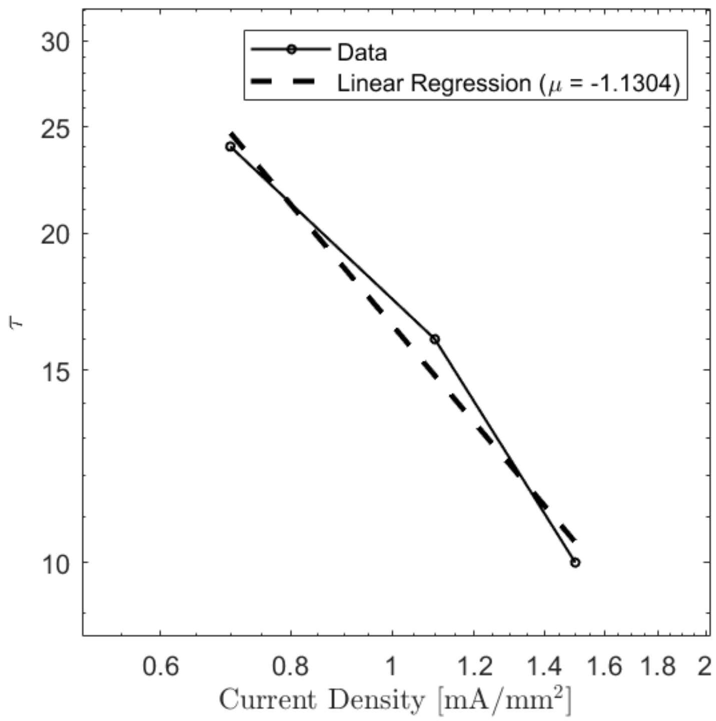


**Supplementary Figure 6** Linear regression of threshold time T as a function of current density, plotted in log-log form. T is defined to be the time at which the normalized area curve for each current density reaches some arbitrary threshold normalized area. The slope of the fit is then used to rescale the normalized area curves onto a single master curve (see Fig. 1C).

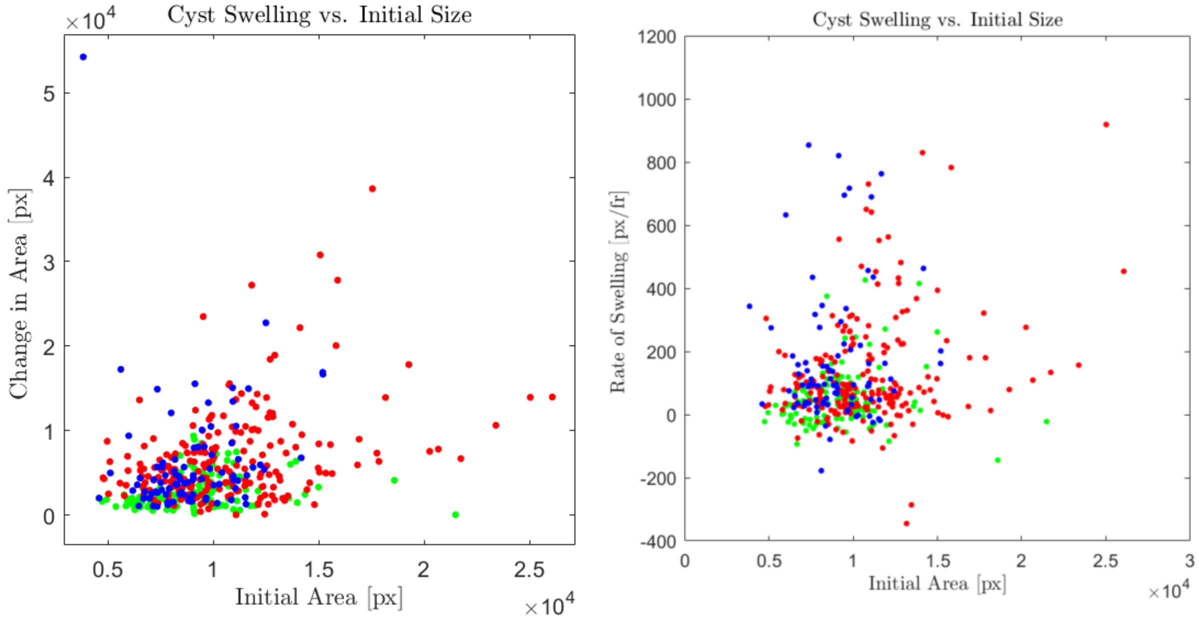


**Supplementary Figure 7 Left:** Maximum normalized area vs. initial area of each cyst after 3 hours of stimulation at low (green), medium (red), and high (blue) current. Initial size has little to no effect on the final size of the cyst. **Right:** Mean rate of swelling in the first hour of stimulation for all cysts across low (green), medium (red), and high (blue) current. Initial cyst area does not affect the rate of swelling within each experimental condition.

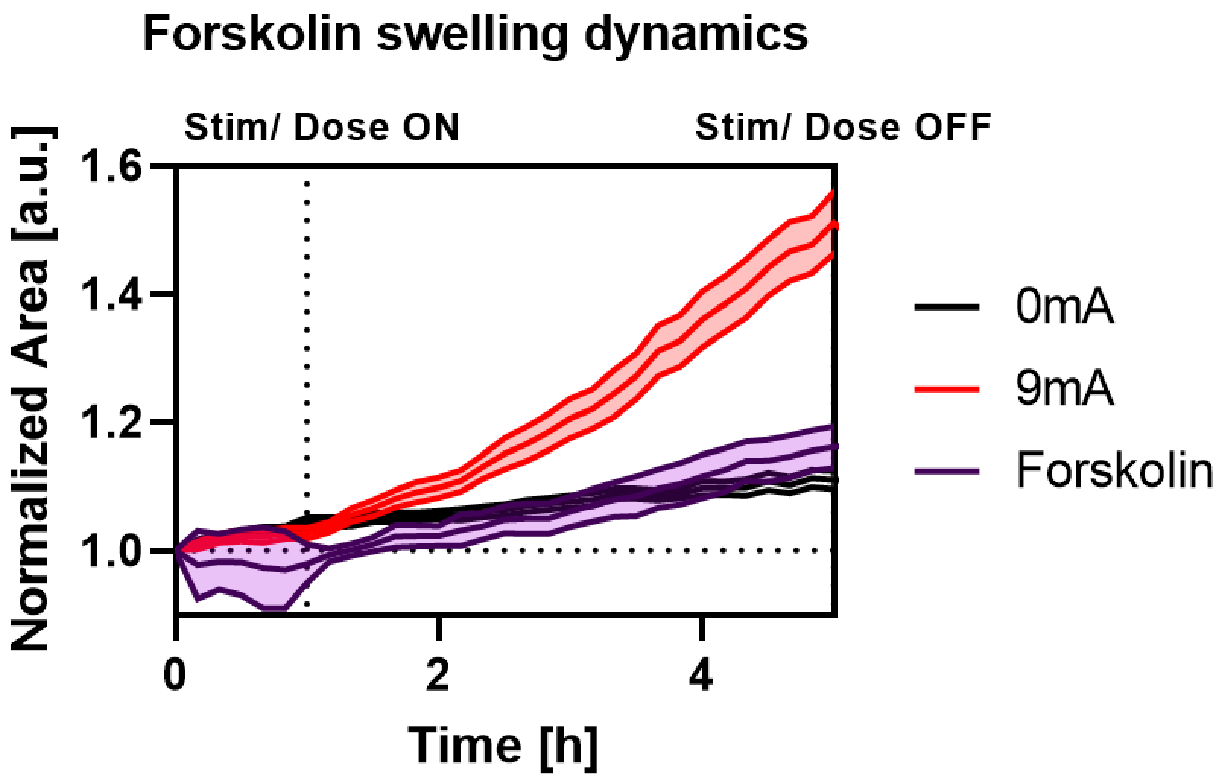


**Supplementary Figure 8** Comparison between 100uM forskolin treatment (purple; N = 1, n = 30) and electrical stimulation at medium current (red; N = 5, n = 150). Electrical stimulation and chemical stimulation is applied at the 1h mark and maintained for 4h.

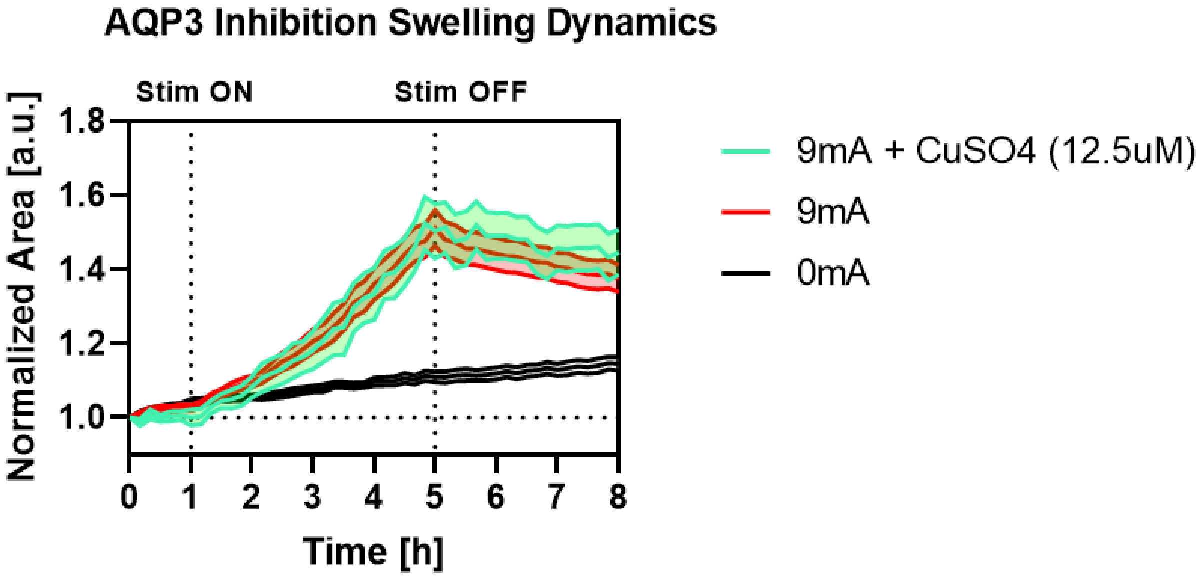


**Supplementary Figure 9** Comparison between MDCK acini without stimulation nor treatment (black; N = 5, n = 145), stimulated with no treatment for 4h (red; N = 5, n = 150), and stimulated with CuSO4 treatment for aquaporin 3 (AQP3) inhibition (teel; N = 3, n = 101). There is no significant reduction in cyst inflation with the inhibition of AQP3.

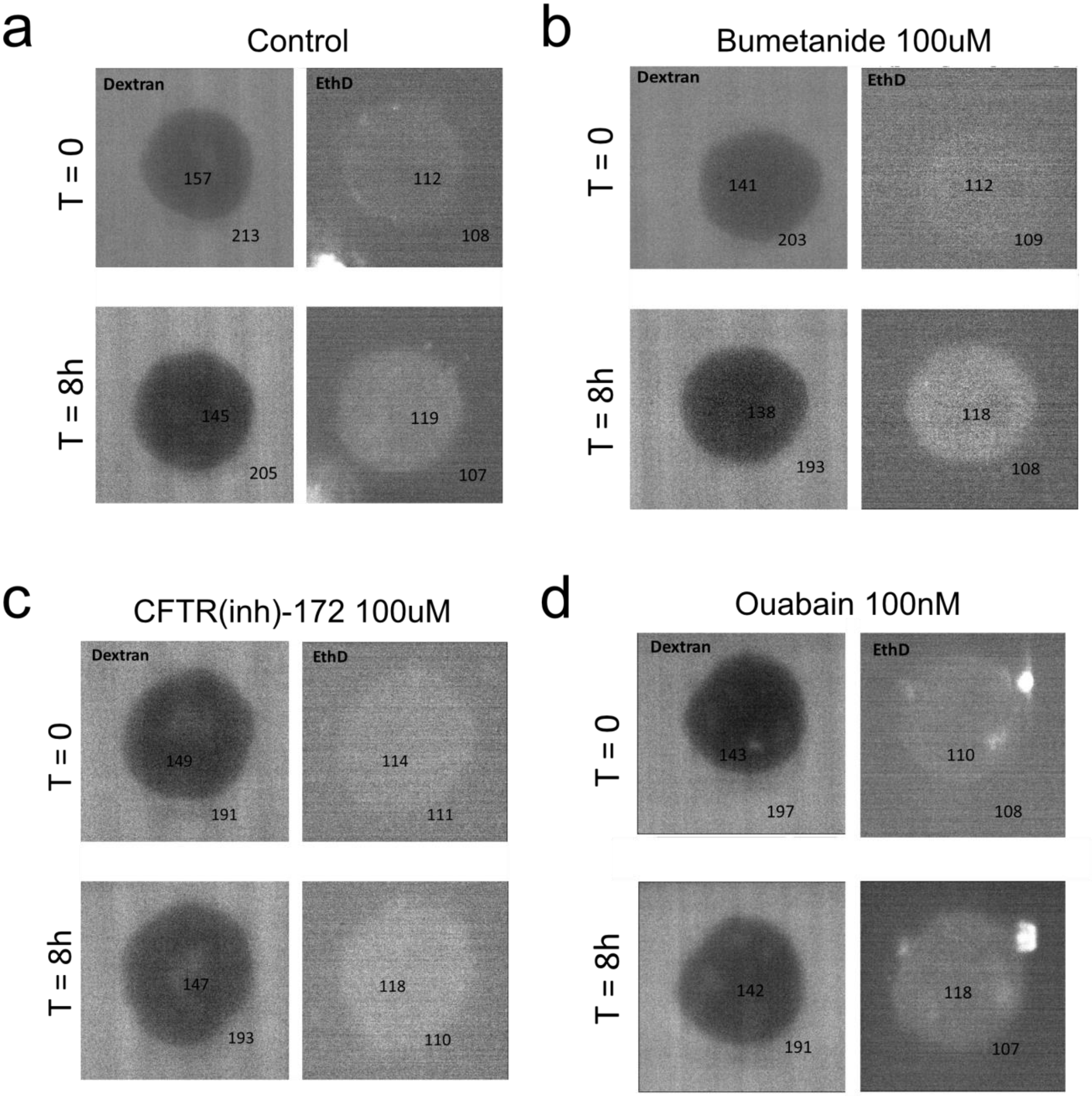


**Supplementary Figure 10**: Dextran and EthD-1 assay to test epithelial barrier function and cytotoxicity with the addition of inhibitors. Inhibitors were added 24h (CFTR(inh)-172, ouabain) or 1h (bumentanide) before the experiment. The fluorescence values within the lumen and external environment are overlayed on the image. In all conditions, the 4kDa dextran did not penetrate the lumen and there was no nuclear localization of EthD-1, confirming that the inhibitor treatment did not induce the concerned effect to our model systems.

| | Cytoskeletal Inhibitors | | | | | |
|---|---|---|---|---|---|---|
| | Control | 1.1 mA/mm$^2$ | Y27632 + 1.1 mA/mm$^2$ | Y27632 | Blebbistatin + 1.1 mA/mm$^2$ | Blebbistatin |
| Control | 1 | | | | | |
| 1.1 mA/mm$^2$ | <0.0001 (****) | 1 | | | | |
| Y27632 + 1.1 mA/mm$^2$ | <0.0001 (****) | <0.0001 (****) | 1 | | | |
| Y27632 | <0.0001 (****) | <0.0001 (****) | <0.0001 (****) | 1 | | |
| Blebbistatin + 1.1 mA/mm$^2$ | <0.0001 (****) | <0.0001 (****) | 0.7247 (n.s) | <0.0001 (****) | 1 | |
| Blebbistatin | <0.0001 (****) | <0.0001 (****) | <0.0001 (****) | 0.0087 (**) | <0.0001 (****) | 1 |

| | Ion Channel Inhibitors | | | | | | | | | |
|---|---|---|---|---|---|---|---|---|---|---|
| | Control | 1.1 mA/mm$^2$ | CFTR-172 50uM + 1.1 mA/mm$^2$ | CFTR-172 100uM + 1.1 mA/mm$^2$ | CFTR-172 100uM | Bumetanide 100uM + 1.1 mA/mm$^2$ | Bumetanide 100uM | Ouabain 100 nM | Ouabain 100 nM + 1.1 mA/mm$^2$ | |
| Control | 1 | | | | | | | | | n.s. |
| 1.1 mA/mm$^2$ | <0.0001 (****) | 1 | | | | | | | | (*) |
| CFTR-172 50uM + 1.1 mA/mm$^2$ | <0.0001 (****) | <0.0001 (****) | 1 | | | | | | | (**) |
| CFTR-172 100uM + 1.1 mA/mm$^2$ | <0.0001 (****) | <0.0001 (****) | <0.0001 (****) | 1 | | | | | | (***) |
| CFTR-172 100uM | <0.0001 (****) | <0.0001 (****) | <0.0001 (****) | <0.0001 (****) | 1 | | | | | (****) |
| Bumetanide 100uM + 1.1 mA/mm$^2$ | <0.0001 (****) | 0.0127 (*) | <0.0001 (****) | <0.0001 (****) | <0.0001 (****) | 1 | | | | |
| Bumetanide 100uM | 0.0036 (**) | <0.0001 (****) | <0.0001 (****) | <0.0001 (****) | <0.0001 (****) | <0.0001 (****) | 1 | | | |
| Ouabain 100 nM | 0.0433 (*) | <0.0001 (****) | <0.0001 (****) | <0.0001 (****) | 0.0172 (*) | <0.0001 (****) | 0.0144 (*) | 1 | | |
| Ouabain 100 nM + 1.1 mA/mm$^2$ | <0.0001 (****) | 0.0201 (*) | 0.8205 (n.s.) | <0.0001 (****) | <0.0001 (****) | 0.0689 (n.s.) | <0.0001 (****) | <0.0001 (****) | 1 | |

**Supplementary Figure 11:** Lookup table of p-values for all experimental conditions. Colors represent significance as measured by a pairwise parametric t-test with Welch's correction. Data distributions were verified to be gaussian by fitting using MATLAB.

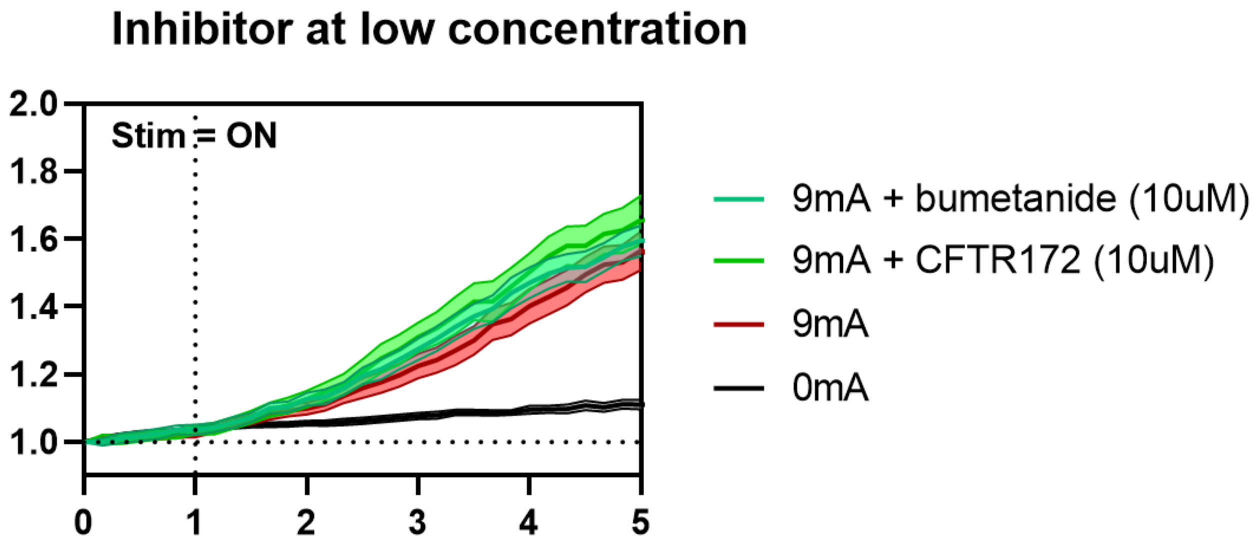


**Supplementary Figure 12**: Swelling dynamics for lower concentration of inhibitors. CFTR(inh)-172, 10uM (green; N = 2, n = 78) and bumetanide 10uM (teel; N = 3, n = 150). There were no significant differences in inflation with the

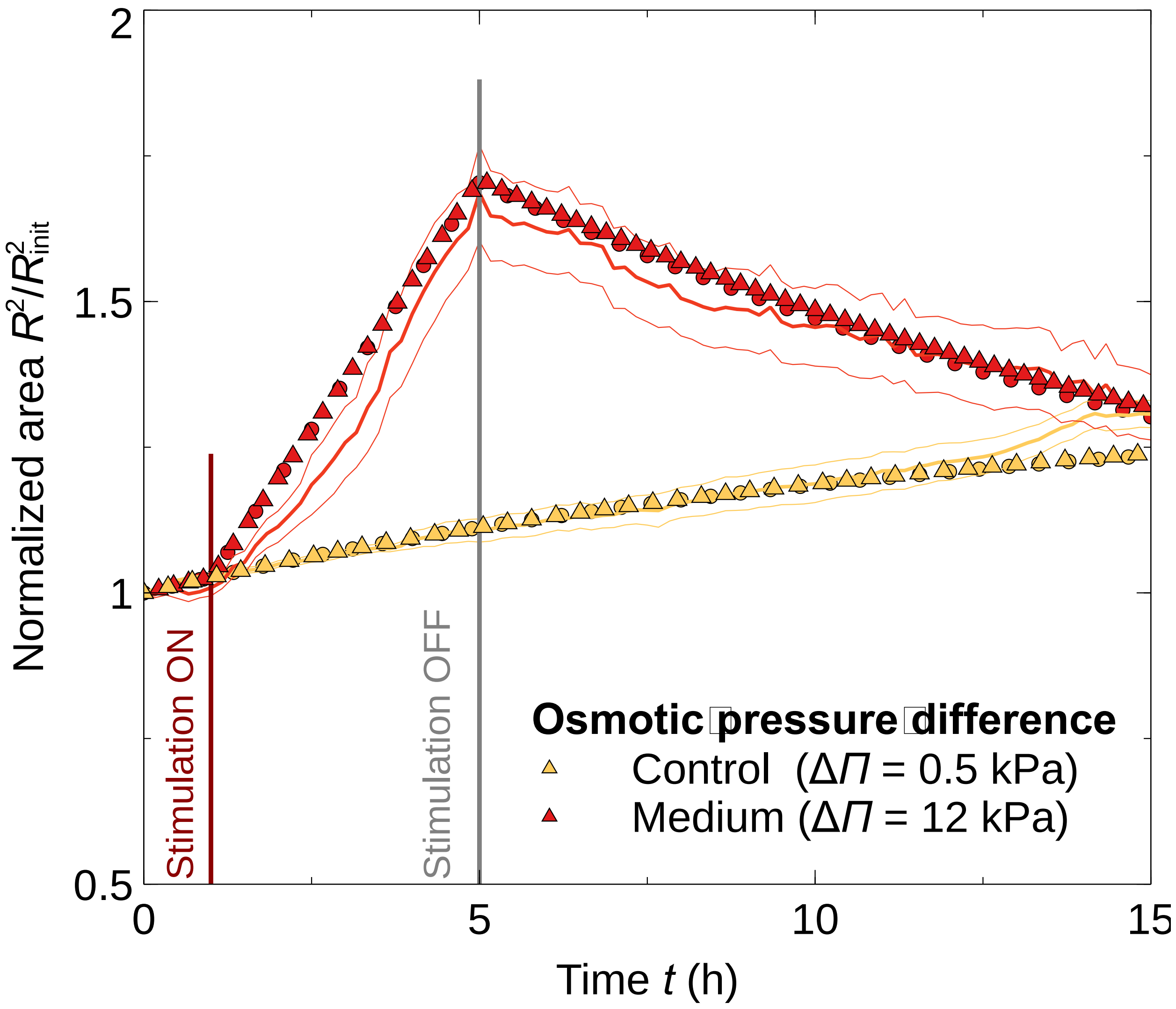


**Supplementary Figure 13** Normalized area as a function of time obtained from experiments (curves) and 1D model (triangles) considering a Maxwell material. In the model, we consider that the osmotic pressure increases drastically due to stimulation but remains constant over time. When the field is turned off, the osmotic pressure is decreased drastically, e.g., due to ion rearrangement or fluid leakage. The build-up hydrostatic normal stress overcomes the osmotic pressure, thus driving cyst shrinkage. See Supplementary Information for further details.

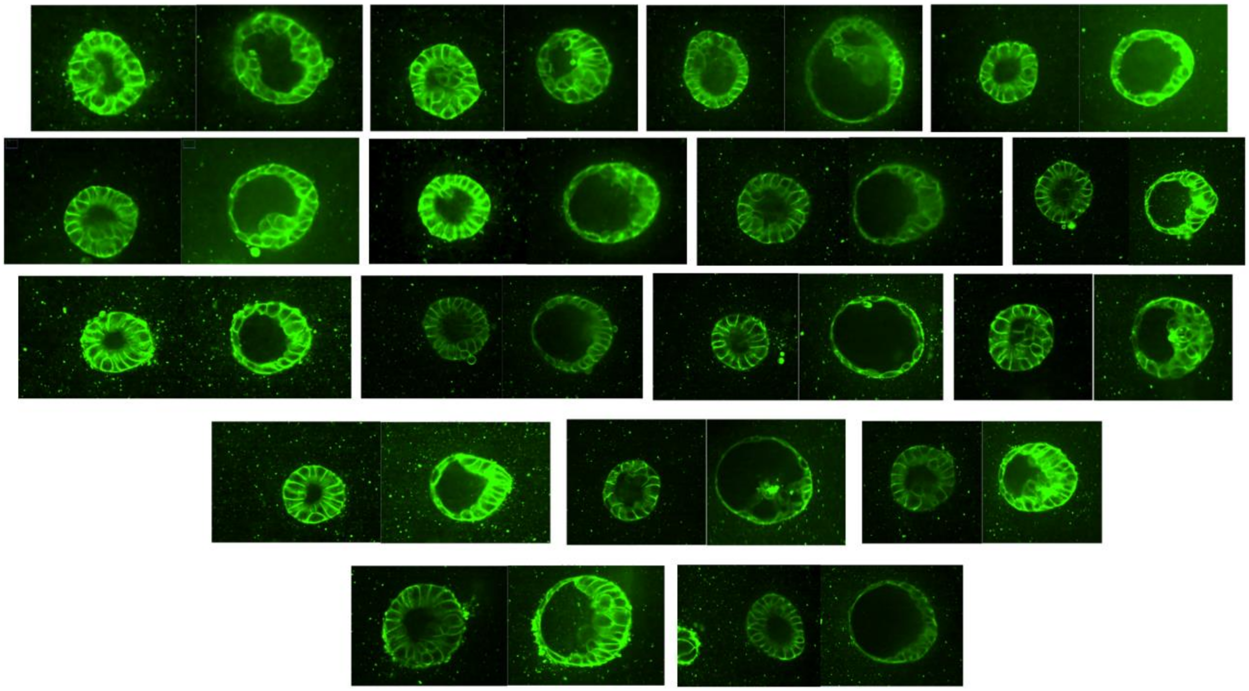

**Supplementary Figure 14** The heterotypic response of electrically stimulated cysts. Some cysts undergo significant areal increase with a radially symmetric response, whereas some cysts do not swell as much, and will instead thicken on the cathode-facing half and thin on the anode-facing half.

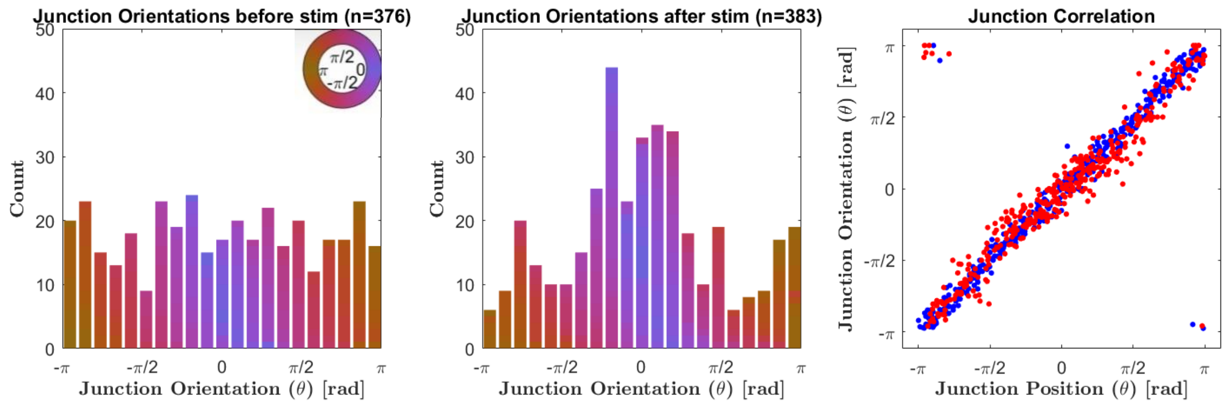


**Supplementary Figure 15 Left**: Uniform distribution of junction orientations before stimulation. **Middle**: Distribution of junction orientations after stimulation shows that junctions on the cathode-facing half of the cyst tend to be more horizontally oriented. **Right**: Scatter plot of junction orientation vs. junction position. Before (blue) and after (red) stimulation, there is a linear relationship between position and orientation.

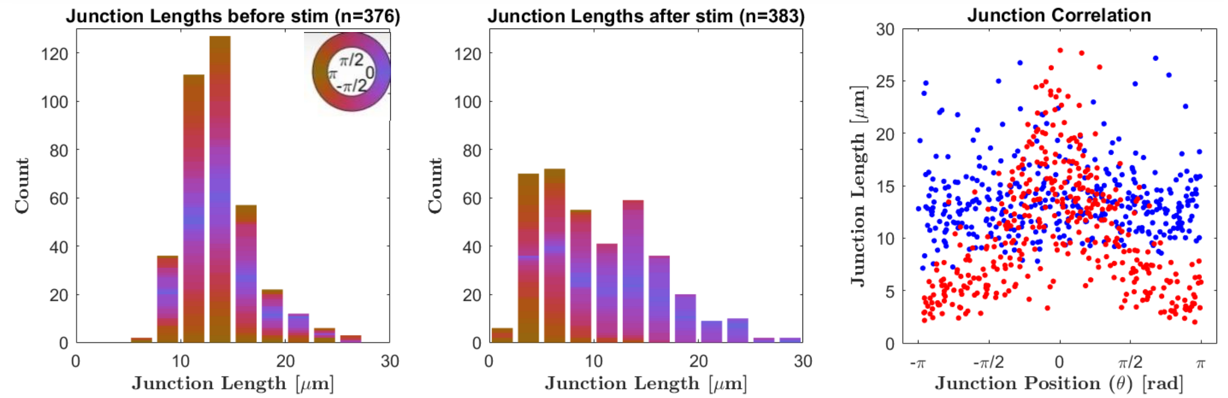


**Supplementary Figure 16 Left:** Distribution of junction lengths before stimulation showing that junctions throughout the perimeter of the cyst tend to have a mean length of ~15um. **Middle:** Distribution of junction lengths after stimulation showing that junctions on the anode-facing half of the cyst tend to be shorter than the original mean length of ~15um whereas junctions on the cathode-facing half of the cyst tend to be longer than ~15um. **Right**: Scatter plot of junction length vs. junction position. Before (blue) stimulation, there is no relationship between junction length and position in the cyst circumference, whereas after (red) stimulation, junctions tend to be longer on the cathode-facing half of the cyst ($-\pi/2 < \theta < \pi/2$).

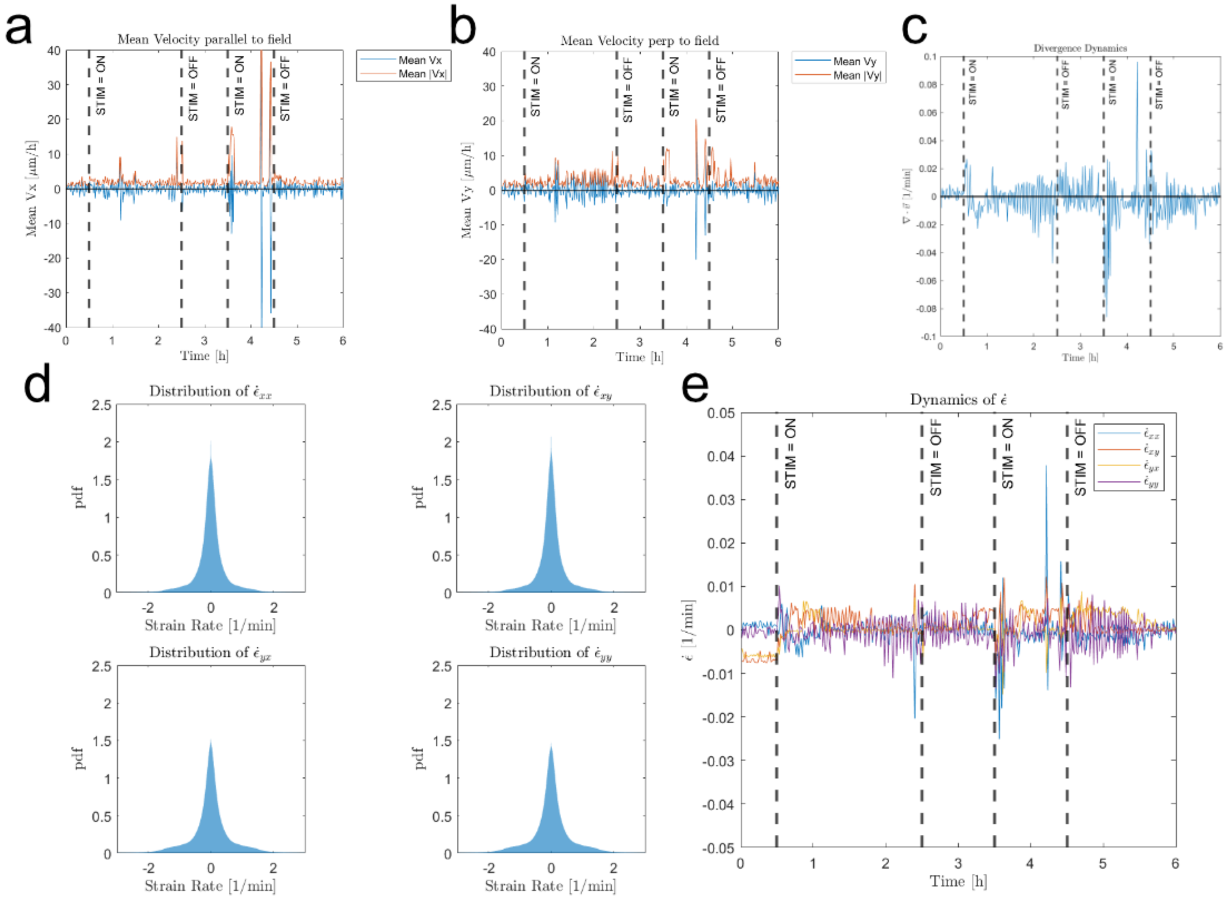


**Supplementary Figure 17** Several deformation analyses were performed on electrically stimulated hydrogels that did not contain any MDCK cysts. (a-b) Mean velocity and speed (a) parallel and (b) perpendicular to the field showed no significant change during stimulation periods. (c) Divergence of the velocity field in the hydrogel remained negligible with and without stimulation. (d) Probability density functions of all entries in the strain rate tensor in the hydrogel during stimulation. Distributions are centered at and symmetrical about zero, suggesting that the gel is not undergoing any directed stress. (e) Temporal dynamics of mean strain rate remain zero during all times, with and without stimulation.

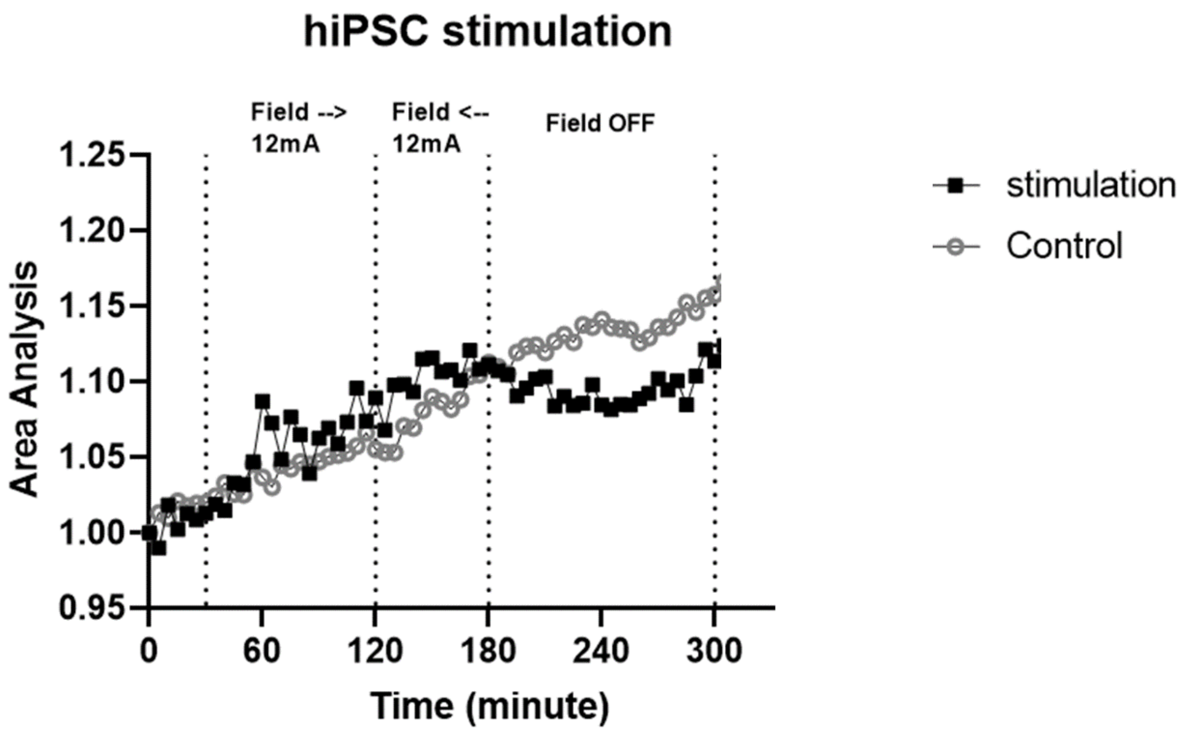


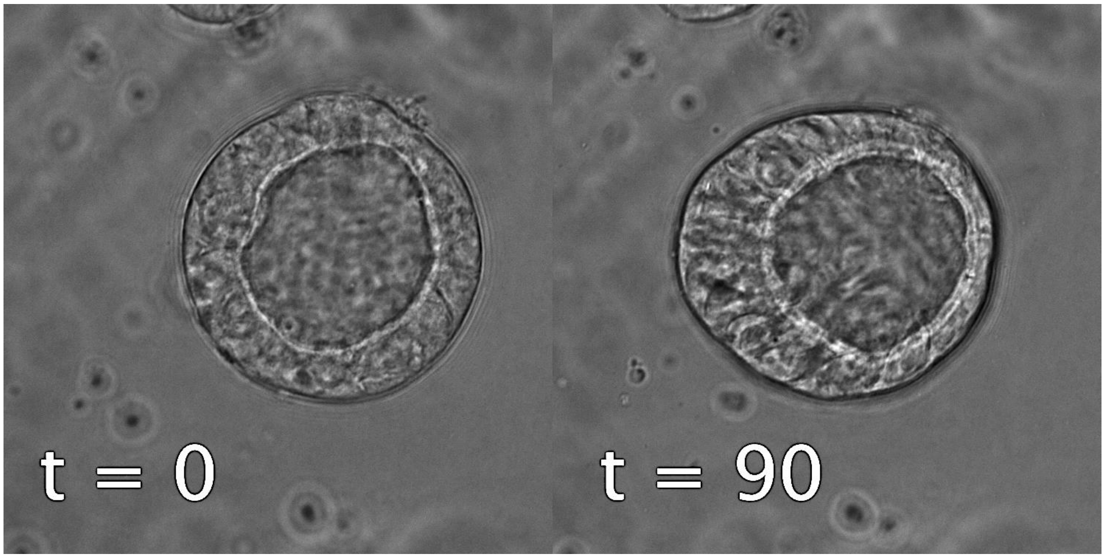


**Supplementary Figure 18** Up: Comparison of inflation dynamics of hiPSC lumenized structures with (stimulation; N =1, n = 12) and without (control; N = 1, n = 47) electrical stimulation. Electrical stimulation does not seem to induce luminal inflation in hiPSC structures unlike in MDCK cysts.

Down: hiPSC spherical structures at initial timepoint (t = 0) and after 90 min of electrical stimulation at 12mA (t = 90 min). The structure does not show luminal inflation but clearly shows thinning-thickening asymmetry. Scale bar = 25um.

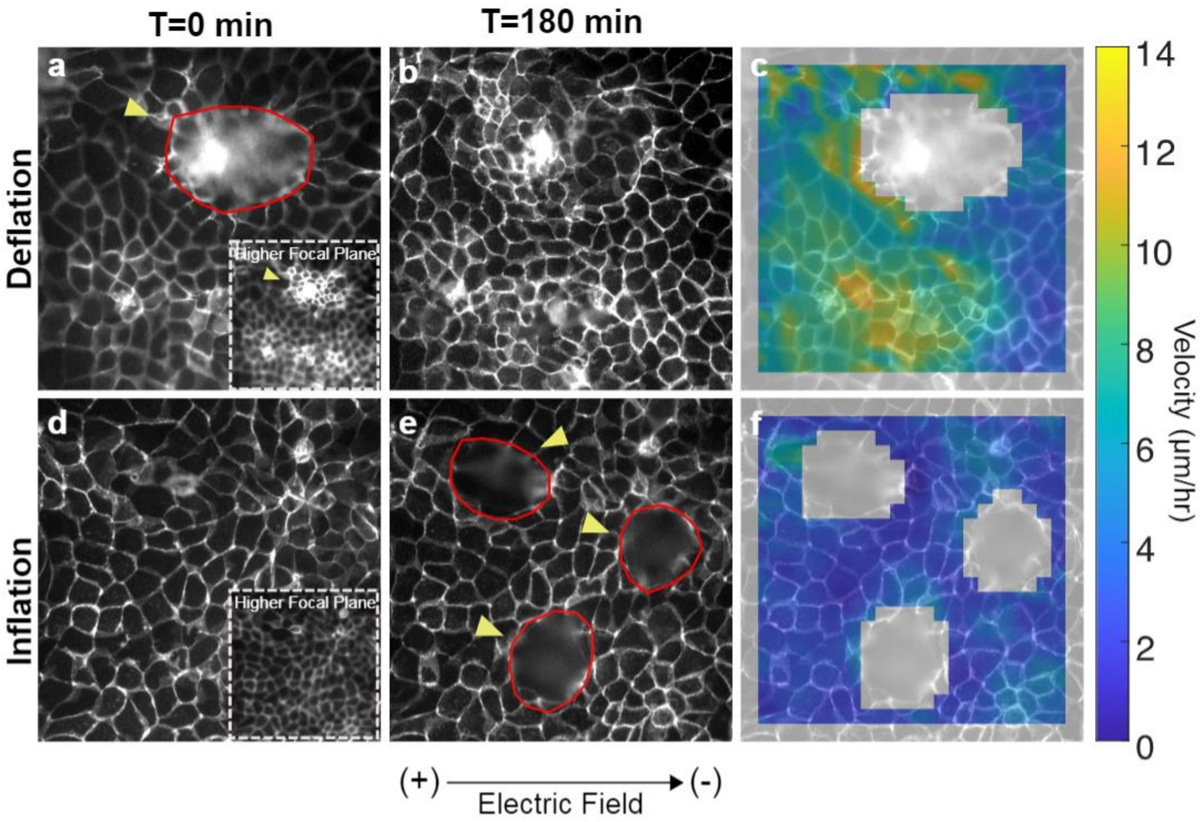


**Supplementary Figure 19**: **a.** MDCK dome (arrow) fully inflated, pre-stimulation. In fluorescence imaging, domes will appear as blurry or dark zones because they are high above the focal plane of the objective. Locally blurred regions of the image mean differences in 3D height (domes). To demonstrate this we show data from different focal planes (see inset). **b.** MDCK dome (arrow) fully deflated, post-stimulation. **c.** Heat map of velocity near the dome that deflates during stimulation, showing high migrational speeds in the region proximal to the dome. **d.** MDCK monolayer (*without* domes) pre-stimulation (see inset showing no in-focus cells out of plane). **e.** MDCK domes (arrows) that have inflated during stimulation. **f.** Heat map of velocities near the inflated domes, showing little to no movement proximal to the domes. Scale bar 25um.

### Supplementary Movies

**Supplementary Video 1** 40x confocal image video of MDCK cyst inflating isotropically under medium current stimulation. Green: membrane dye (Memglow 488); cyan: nuclei (Hoechst 33342). Electrical stimulation indicated with "Field →". 3h stimulation on. 10 min/frame. Scale bar = 50um.

**Supplementary Video 2** 40x confocal image video of MDCK cyst undergoing structural asymmetry as it inflates under medium current stimulation. Green: membrane dye (Memglow 488); cyan: nuclei (Hoechst 33342). Electrical stimulation indicated with "Field →". 3h stimulation on. 10 min/frame. Scale bar = 50um.

**Supplementary Video 3** 10x transmitted light image video and binarization of inflating MDCK cyst under medium current stimulation. Electrical stimulation indicated with "Field →". 1h stimulation off, 4h stimulation on, 3h stimulation off. 10 min/frame. Scale bar = 50um.

**Supplementary Video 4** 10x transmitted light (left; grey) and EthD-1 live imaging (right; red) for cellular viability check. Structure inflates steadily with electrical stimulation. Electrical stimulation indicated with "Field →". 1h stimulation off, 4h stimulation on, 3h stimulation off. 10 min/frame. Scale bar = 50um.

**Supplementary Video 5** 10x transmitted light (left; grey) and EthD-1 live imaging (right; red) for cellular viability check. Structure inflates rapidly, ruptures, and deflates in the presence of electrical stimulation. Electrical stimulation indicated with "Field →". 1h stimulation off, 4h stimulation on, 3h stimulation off. 10 min/frame. Scale bar = 50um.

**Supplementary Video 6** 40x confocal image video of MDCK cyst treated with 100uM blebbistatin inflating under medium current stimulation. Green: membrane dye (Memglow 488); cyan: nuclei (Hoechst 33342). Electrical stimulation indicated with "Field →". 2.5min/ frame. Scale bar = 50um.

**Supplementary Video 7** 40x confocal image video of inflating RFP-Ecadherin MDCK cyst under oscillating directional field at 1.1mA/mm^2 stimulation. Direction of electrical stimulation indicated with "Field →" and "Field ←". 15 min control, 1h stimulation with field going to the right, 1h stimulation with field going to the left, 1h stimulation off. 15 min/frame. Scale bar = 50um.

**Supplementary Video 8** 40x confocal image video of MDCK cyst undergoing structural asymmetry but exhibiting minimal luminal inflation under medium current stimulation, suggesting inflation and asymmetry are uncoupled. Green: membrane dye (Memglow 488); cyan: nuclei (Hoechst 33342). Electrical stimulation indicated with "Field →". 3h stimulation on. 10 min/frame. Scale bar = 50um.

**Supplementary Video 9** 40x confocal image video of lumenized hiPSC structures forming asymmetry under high current stimulation (12mA). Green: GFP-actin; cyan: nuclei (Hoechst 33342). Electrical stimulation indicated with "Field →". 2.5min/ frame. Scale bar = 50um.

**Supplementary Video 10** 40x confocal image video of MDCK cyst inflating under electrical stimulation. Individual nuclei were tracked with TrackMate function in ImageJ to confirm that cells migrated along the electrical field vector lines that wrap around the cyst as shown in computational results. Migration tracks are temporally color-coded and proceed along the parula colormap on the lower left (blue -> red). Electrical stimulation indicated with "Field →". 10min/ frame. Scale bar = 20um.

**Supplementary Video 11** 5x transmitted light image video of MDCK monolayers treated with 50uM LY294002 migrating towards the anode (non-treated monolayers migrate towards the cathode). Electrical stimulation indicated with "Field →". 10min/ frame. Scale bar = 500um.

**Supplementary Video 12** 40x confocal image video of MDCK cyst treated with 50uM LY294002 inflating under medium current stimulation. Fluorescence: membrane (MemGlow 488). Electrical stimulation indicated with "Field →". 10min/ frame. Scale bar = 20um.

**Supplementary Video 13** 20x epifluorescence image video of RFP-E-cadherin MDCK domes with no treatment deflating with electrical stimulation. Note the high migrational speeds near the dome. Field directed towards the right, 9mA. 20 min/frame. Scale bar = 100um.

**Supplementary Video 14** 20x epifluorescence1 image video of RFP-E-cadherin MDCK domes treated with 25uM LY294002 inflating with electrical stimulation. Note the low migrational speeds near the dome. Field directed towards the right, 9mA. 20 min/frame. Scale bar = 100um.

**Supplementary Video 15** 20x transmitted light image video of inflating mouse intestinal stem cell organoid under medium current stimulation. Left crypt inflates significantly with stimulation. Electrical stimulation indicated with "Field →". 30 min stimulation off, 2h 30 min stimulation on, 1h stimulation off. 1 min/frame. Scale bar = 100um.

**Supplementary Video 16** 20x transmitted light image video of inflating mouse intestinal stem cell organoid under medium current stimulation. Left crypt inflates significantly with stimulation. Electrical stimulation indicated with "Field →". 30 min stimulation off, 2h 30 min stimulation on, 1h stimulation off. 1 min/frame. Scale bar = 100um.
